# Protocol for genotyping cephalopod sex using a skin swab and quantitative PCR

**DOI:** 10.64898/2026.03.31.715692

**Authors:** Frederick A. Rubino, Connor J. Gibbons, Thomas J. Mungioli, Scott T. Small, Gabrielle C. Coffing, Andrew D. Kern, Tessa G. Montague

## Abstract

The coleoid cephalopods (octopus, cuttlefish, and squid) are emerging model organisms for neuroscience, development, and evolutionary biology. Determining their sex early in life is critical for population management and controlled experiments. Here, we present a protocol to non-invasively determine the sex of multiple cephalopod species as young as 3 hours post-hatching using a skin swab and quantitative PCR (qPCR). We describe steps for designing qPCR primers, swabbing live animals, extracting DNA, running the qPCR, and analyzing the results. For complete details on the use and execution of this protocol, please refer to Rubino et al.^1^

**Highlights:**

- Swab live cephalopods as early as 3 hours post-hatching
- Extract DNA from cephalopod skin swabs
- Perform qPCR-based sex determination
- Design and validate qPCR primers for new species

**Graphical abstract:** 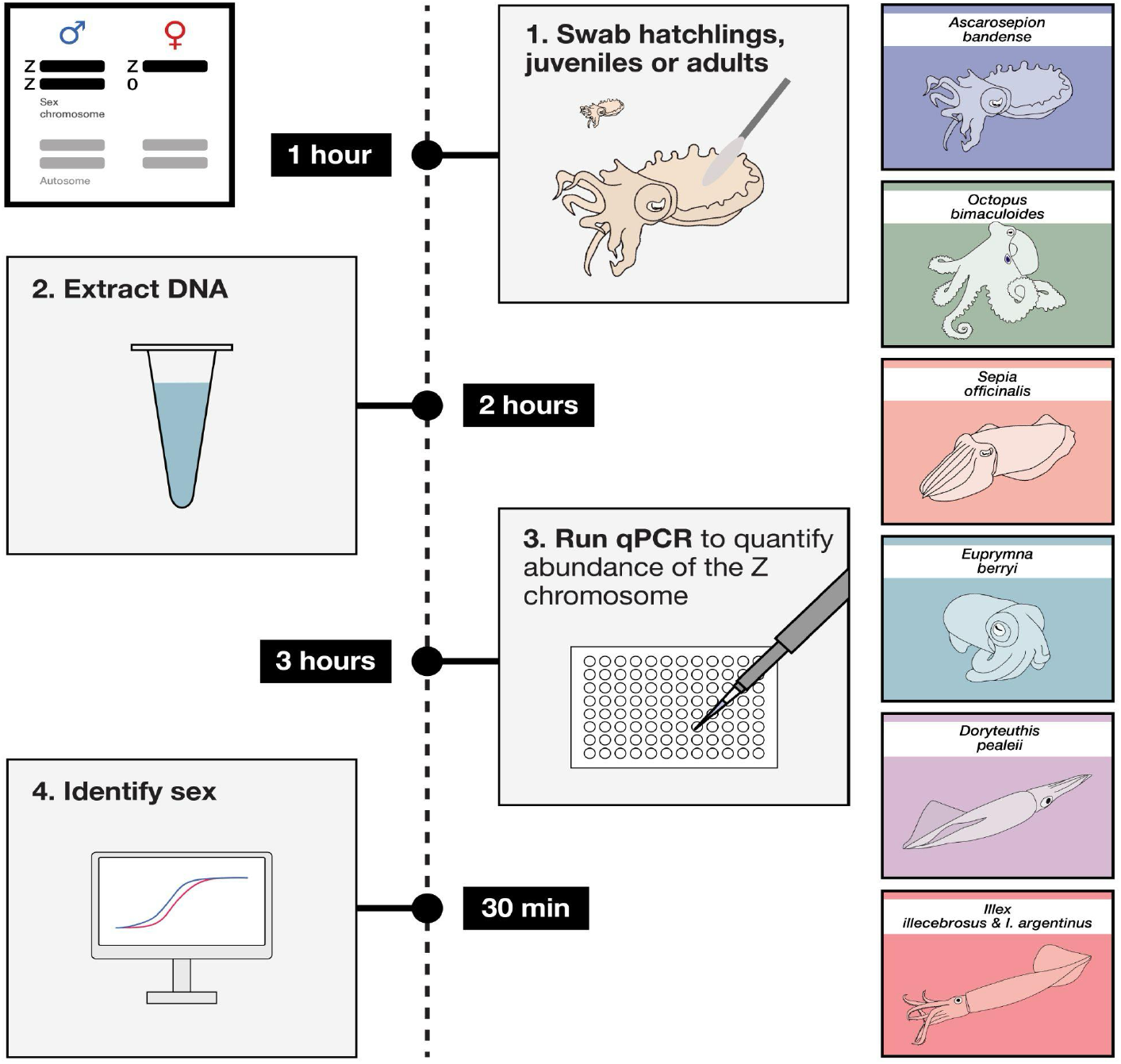

## Before you begin

Coleoid cephalopods (octopus, cuttlefish, and squid) are highly derived mollusks with the largest brain-to-body ratios among invertebrates and an array of complex behaviors, including adaptive camouflage, problem-solving, episodic-like memory, and dynamic social behaviors^2-5^. Cephalopods diverged from vertebrates over 600 million years ago^6^, yet convergently evolved large, complex brains, camera-type eyes, and closed circulatory systems^7^. Thus, the coleoid cephalopods combine unique biological features – including extensive RNA editing^8^, an elaborate central and distributed nervous system^9^, and adaptive camouflage^3^ – with a powerful comparative system for uncovering general principles of nervous system organization, behavior, and evolution.

Historically, cephalopod research has relied on wild-caught species such as the longfin inshore squid (*Doryteuthis pealeii*), which played a central role in the discovery of the action potential^10^. Advances in aquaculture have enabled the culture of multiple species in the laboratory, including the California two-spot octopus (*Octopus bimaculoides*), common cuttlefish (*Sepia officinalis*), dwarf cuttlefish (*Ascarosepion bandense*), and hummingbird bobtail squid (*Euprymna berryi*), driving rapid expansion of cephalopods as experimental systems.

A key limitation in cephalopod culture is the difficulty of establishing breeding populations with defined sex ratios at early developmental stages, which can strongly influence egg production and aggression^1^. Sex can be determined in adults by the presence of sexually dimorphic traits, such as a hectocotylus (a modified arm found in males of some cephalopod species), but these traits can be difficult or impossible to identify in juveniles. Early sex identification is therefore critical for selective breeding, including preventing unintended sperm storage, as well as for behavioral studies, where some species exhibit sex mimicry^11,12^. The ability to determine sex in wild-caught fishery species, such as the northern shortfin squid (*Illex illecebrosus*) and Argentine shortfin squid (*Illex argentinus*), could also enable improved population monitoring and fisheries management.

The recent identification of a genetic sex determination system in octopus^13^ revealed a ZZ sex karyotype in males and Z0 in females. This system of sex determination can be detected via qPCR by measuring the ratio of an autosomal amplicon to a Z-linked amplicon, yielding an expected 1:0.5 ratio in females and 1:1 ratio in males^1^. Here, we describe a non-invasive protocol for sex determination in cephalopods using skin swabs and qPCR, which can be performed on live animals as young as 3 hours post-hatching. We provide validated primer sets for *O. bimaculoides, S. officinalis, A. bandense, E. berryi, D. pealeii, I. illecebrosus*, and *I. argentinus*, along with guidelines for designing primers for additional species.

### Optional: Primer design and validation

This protocol includes validated primer sequences for seven cephalopod species (Graphical Abstract). To adapt this protocol for an additional cephalopod species, this section provides instructions to design stringent qPCR primers and validate them. Note that some DNA sequence data are required, although a fully assembled genome is not necessary^1^. To facilitate identification of candidate autosomal and Z-linked targets, **Table S1** provides amino acid sequences for four autosomal genes and four Z-linked genes from eight cephalopod species that can be used to identify equivalent sequences in your species of interest.

**CRITICAL:** Primers must be designed to amplify genomic DNA, not cDNA or mRNA-derived sequence, because primers separated by introns may fail to amplify the intended genomic target. When only RNA-seq data are available, design multiple candidate primer pairs and screen them for amplification of a product of the expected size.

#### Primer design

*Timing: 2 hours*

**Note:** This protocol uses Geneious Prime (which uses a modified version of Primer3 2.3.7) for primer design; however, equivalent primer design software can be used with the same design criteria.

1. Import genomic DNA sequences corresponding to candidate autosomal loci into Geneious using the “Import from File” or database retrieval functions. Each locus should be 15-30 kb in length.
  a. Include several dozen loci to increase the likelihood of identifying suitable primer pairs. In practice, ∼24 loci are usually sufficient to give 12 good primer pair candidates, but it is easier to include more loci and design an excess of primer pairs at this step.
  b. When possible, select loci distributed across the chromosome to minimize regional bias.
2. Use the Primer Design tool to generate a candidate qPCR primer pair against each locus with the following parameters:
  a. Set primer pair penalty weights high (e.g. 5) for individual and pair “dimer Tm” parameters under “Primer Picking Weights” in Geneious Prime to minimize primer–dimer formation.
3. Select primer pairs that lack predicted secondary structure, primer-dimer formation, or significant off-target binding within the available sequence dataset.
  a. Screen candidate primer pairs using OligoAnalyzer Tool for hairpins, self-dimers, and hetero-dimers. Reject primer pairs with predicted secondary structures with a Tm >30°C or with ≥4 consecutive complementary bases in primer–dimers, particularly at the 3^′^ end.
  b. Perform a BLAST search of each predicted amplicon against the reference genome (or available sequencing data) to assess off-target binding. Reject primers that bind repetitive regions and could generate similarly sized off-target products.
4. Repeat steps 1–3 for candidate sex-linked (Z-linked) loci.
5. Design a minimum of 12 primer pairs for the autosomal loci and 12 primer pairs for the Z-linked loci (24 total primer pairs recommended).

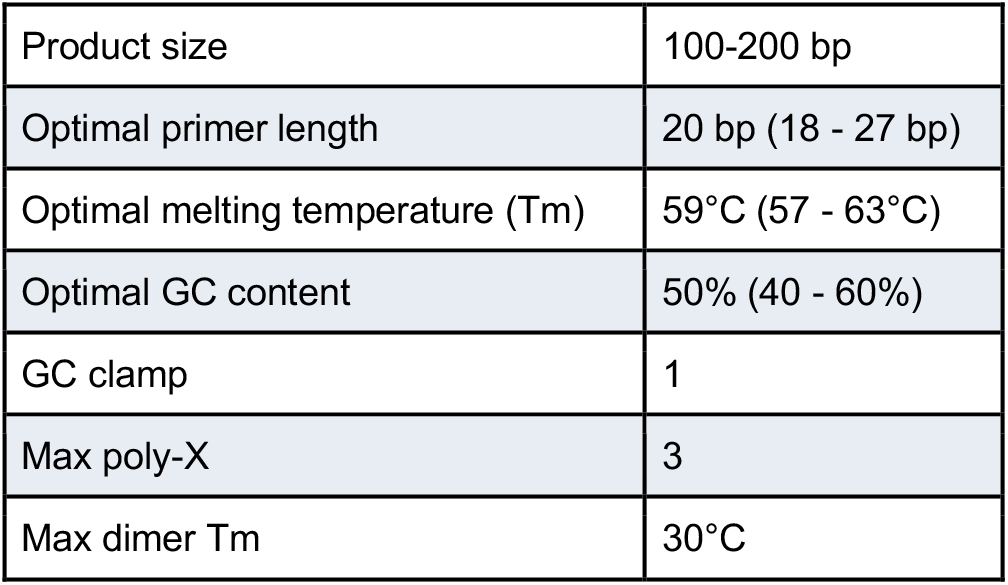

|  |  |
| --- | --- |
| Product size | 100-200 bp |
| Optimal primer length | 20 bp (18 - 27 bp) |
| Optimal melting temperature (T<sub>m</sub>) | 59°C (57 - 63°C) |
| Optimal GC content | 50% (40 - 60%) |
| GC clamp | 1 |
| Max poly-X | 3 |
| Max dimer T<sub>m</sub> | 30°C |

#### Primer validation

*Timing: 6-8 hours (excluding primer synthesis)*

1. Obtain genomic DNA (gDNA) from confirmed male and female individuals of the target species for validation.
  a. For instance, extract gDNA from post-mortem tissue (e.g. arm tips) of phenotypically sexed animals by following the manufacturer’s instructions of the Monarch Spin gDNA Extraction Kit. This should yield sufficient DNA for multiple rounds of primer testing.
2. Make a 20× stock primer mix in nuclease-free water – 5 µM forward & 5 µM reverse primer.
3. Set up qPCR reactions for each primer pair using gDNA from one confirmed male and one confirmed female individual. Include a no-template control (NTC) for each primer pair (see **“qPCR amplification of autosome and sex chromosome loci”** for complete details for qPCR reactions). For each reaction:
  a. Use 2 µL of 1 ng/µL gDNA as template.
  b. Use primers at a final concentration of 0.25 µM.
  c. Perform 4 technical replicates per condition and calculate the mean Cq value.
4. Evaluate amplification curves and discard primer pairs that show:
  a. No amplification
  b. Late amplification (e.g. Cq > 35)
  c. Amplification in NTCs
  d. Early amplification.
    i. Most primer pairs should give Cq values that cluster within a small range (∼2 Cq). Early Cq values likely indicate off-target binding or primer-dimer amplification.
5. Examine melt curves and discard primer pairs that produce multiple peaks or evidence of non-specific amplification.
  a. Accept only primer pairs with single, sharp peaks that are consistent between male and female samples and absent in NTCs.
6. Resolve PCR products on a 2% agarose gel to confirm a single amplicon of the expected size using a 100 bp ladder.
7. Confirm amplicon identity by sequencing, either Sanger sequencing (forward and reverse reactions) or using a direct sequencing approach (e.g. nanopore “Premium PCR Sequencing” service by Plasmidsaurus).
8. For primer pairs that pass validation (steps 1-5), compare Cq values between male and female samples:
  a. Compute ΔCq(Auto - Sex) for each primer pair in both sexes. **Note:** Given that males have two copies of the Z chromosome (ZZ), whereas females have one (Z0), a two-fold difference in template corresponds to a ∼1 cycle difference in Cq.
  b. Identify combinations of autosomal and sex-linked primer pairs that satisfy: ΔCq (Auto − Sex, male) − ΔCq (Auto − Sex, female) ≈ +1.0
  c. Accept combinations of primer pairs with values between 0.8 and 1.2 for further analysis.
  d. Select ≥4 primer pairs per chromosome type that meet these criteria. Individual primer pairs are expected to perform well in multiple autosome–sex primer combinations.
9. Determine amplification efficiency for all primer pairs that passed the validation and selection criteria above. This will inform what concentration ranges the assay will work within:
  a. Prepare a gDNA standard curve from 10 ng/μL to 0.001 ng/µL in an 8-well strip using 3.126× (√(10)×) serial dilutions in nuclease-free water.
  b. Run qPCR reactions as described above (step 4) and record Cq values for each template concentration.
  c. Plot the standard curve as Cq versus log10(template concentration).
    i. **Note:** Standard curves should be linear over approximately Cq 20 - 30.
    ii. Identify the linear range for each primer pair and record the corresponding Cq values. The assay will make reliable predictions only within this range for this primer pair.
  d. Fit a linear regression within the linear range: Cq = m × log10([template]) + b
  e. Calculate primer efficiency: E = 10^(−1/m) − 1 **Note:** (i) 100% efficiency corresponds to slope = −3.32. (ii) Acceptable range: 90–110% efficiency (slopes −3.6 to −3.1).
  f. Discard primer pairs with efficiencies outside this range and retain those with the highest performance.
10. Using the best-performing primer pairs, perform sex determination on 8 animals (ideally 4 female and 4 male) using gDNA normalized to 1 ng/µL:
  a. See section “**qPCR amplification of autosome and sex chromosome loci**” for complete details on qPCR reactions and analysis.
  b. Randomize gDNA samples so that sample identity (sex) is blinded during qPCR setup and analysis.
  c. Follow steps 2-5 to obtain ΔCq(A-S) for all 8 animals, for all primer pair combinations.
  d. Ensure that Cq values fall within the validated linear range for each primer pair (step 6).
    i. If not, adjust template concentrations by dilution or concentration (a two-fold dilution will increase the Cq value by ∼1).
  e. For each primer pair combination (autosome & sex chromosome) assign sex to all 8 samples based on ΔCq values, following the steps in section “**qPCR analysis**”.
11. Identify primer pair combinations that correctly classify all samples (100% accuracy); these are validated for future genotyping.
  a. Maintain a record of genotyping performance for all future trials and note any failures for each primer pair combination.
  b. Depending on the error tolerance of your application, you may want to perform more extensive trials at this point.

**Note:** If no primer pairs pass validation, redesign additional candidates or consider alternative loci.

#### Optional: Build a housing tray for temporary individual isolation

*Timing: ∼6 hours*

To maintain the identity of individual cephalopods after skin swabbing and before sex assignment results, construct a compact housing tray that enables temporary (24-72 h) high-density isolation of up to 20 cephalopods within a single 20” x 15” footprint. Because juvenile and subadult cephalopods can pass through extremely small openings, construct the housing system with tight tolerances and secure plumbing.

#### Construction of the housing tray

1. Laser-cut divider walls from 1/4” cast acrylic.
  a. Use the provided SVG file (**Supplementary File 1**) to cut:
    i. 3 long divider walls (spanning the full length of the tray, front to back, **Figure 1A**).
    ii. 5 short divider walls (spanning left to right between long dividers, **Figure 1A**).
    iii. Use laser cutting settings optimized for 1/4” cast acrylic, adjusting power, speed, and passes according to the specific laser cutter being used.
  b. Ensure slot tolerances are tight enough to prevent lateral movement once inserted into the divider box.
    i. To test fit:
      1. Insert a divider fully into the molded slots of the empty tray (See **Figure 1B** for slot locations).
      2. Attempt to gently shift the divider side-to-side.
    ii. Fit is acceptable if the divider remains upright with minimal lateral movement, and the divider does not shift when the tray is tapped.
    iii. If dividers are too loose, re-cut with slightly increased length (+0.1–0.2 mm).

**Figure 1.**
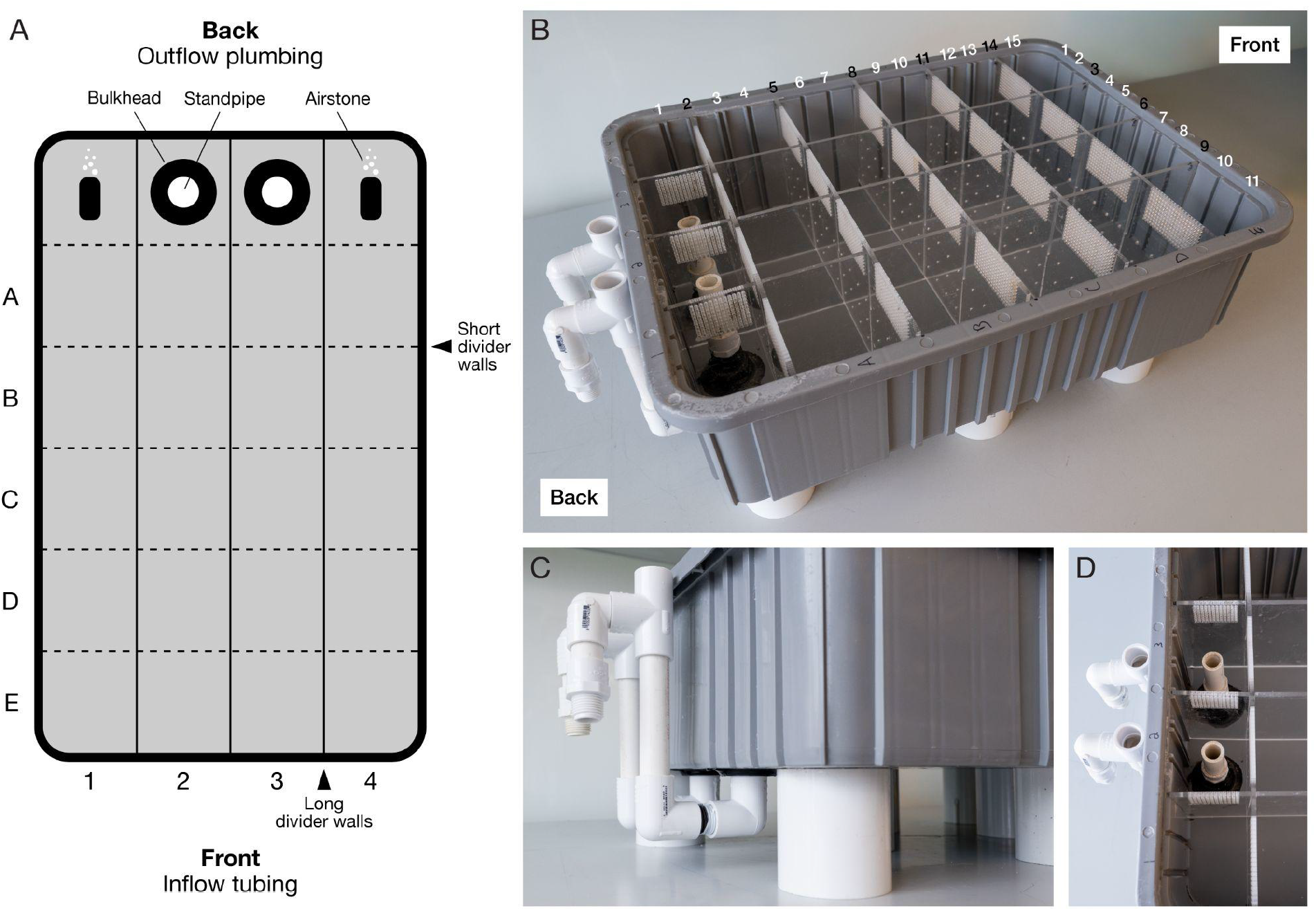
High-density housing tray for temporary isolation of animals. A) Schematic of housing tray containing 20 usable compartments for individual animals. The footprint is compatible with most aquatic rack systems. B) Assembled housing tray. Interlocking ¼” cast acrylic dividers fit tightly into molded slots to prevent animal movement between compartments. C) External bulkhead and PVC standpipe assembly used to establish controlled overflow and consistent water level. D) Internal view of bulkhead placement and vertical standpipes that set the operating water level. **CRITICAL:** Dividers must fit snugly into the tray’s molded slots. Even small gaps can permit hatchlings or juveniles to squeeze between compartments. **Note:** If a laser cutter is not available, acrylic components can be custom ordered from commercial laser cutting services.
2. Prepare outflow plumbing with bulkhead installation.
  a. Identify the back of the tray as the side that will be closest to the sump or plumbed structure that will send the outflowing system water directly to the sump for filtration (**Figure 1A**).
  b. Using a handheld electric drill with a holesaw bit, drill two ⅞” holes in the bottom of the back-most center compartments (**Figure 1A**).
    i. Center each hole within its compartment floor.
    ii. Maintain at least 0.5” clearance from divider walls to ensure proper bulkhead sealing (**Figure 1D**).
  c. Install the threaded bulkhead such that the flush flange sits on the exterior of the tray and the threaded portion protrudes into the interior (**Figure 1**).
    i. Place the gasket on the water-facing (internal) side.
    **Note:** The bulkheads are installed in a non-standard orientation due to space constraints of the existing aquatic system. Reversing the orientation of the threaded side is acceptable, and may be preferred if space permits, provided the gasket remains seated on the water-facing side.
3. Install outflow plumbing.
  a. Apply Teflon tape to all threaded components before installation:
    i. Wrap clockwise (in the direction of tightening).
    ii. Use at least one overlapping layer.
  b. Attach a ½” PVC male threaded adapter to each bulkhead from inside the tray (**Figure 1A**).
  c. Cut and insert the PVC pipe into the male adapters to set the operating water level. The height of the internal standpipe determines the tray water level. A PVC length of 3 ¾” will create a water depth of approximately 5”.
  d. Attach a 90° threaded elbow fitting to each bulkhead on the external side using a ½” threaded nipple (**Figure 1C**):
    i. Orient elbows to direct flow toward the external drain or outflow line returning to the sump.
      1. If utilizing a conventional aquatic rack system where the outflow plumbing sits at or above the bulkhead height, account for the vertical difference by continuing to step 3e.
      2. If the outflow plumbing sits below the bulkhead height, plumb a direct downwards path and proceed to step 4.
  e. Create a U-shaped section of plumbing utilizing a 90° elbow, and straight piece of ½” PVC to reach the required height of your specific aquatic rack system (**Figure 1C**).
    a. Connect the 90° threaded elbows utilizing ½” threaded nipples.
  f. Create a hook utilizing a ½” tee fitting and slip 90° elbow to direct water outflow into the overflow compartment of the aquatic rack system (**Figure 1C**).
    a. The tee fitting is utilized to break any siphon and prevent issues with air locking in the outflow plumbing. **Note:** The installation of a “U” and “hook” structure in the outflow plumbing is required to accommodate constraints of the existing aquatic system and maintain proper water outflow to the sump. Where possible, a simpler configuration – such as attaching a ½” 90° PVC barb to the bulkhead and routing aquarium-safe tubing directly to the sump or filtration area – may be more user-friendly.
4. Insert divider walls.
  a. Place long dividers into vertical slots 3, 6, and 9 (left to right, **Figure 1B**).
  b. Place short dividers into horizontal slots 2, 5, 8, 11, and 14 (front to back, **Figure 1B**). The completed layout creates 20 isolated compartments for animals plus 4 compartments for the outflow and air.
5. Stabilize and level the tray.
  a. Elevate the tray using PVC couplings as risers beneath the base.
    i. Risers are placed in consistent intervals along the outside perimeter and middle of the tray to prevent the bottom of the tray from bowing.
    ii. This elevation provides clearance for outflow plumbing and ensures compatibility with aquatic rack systems that use elevated outflow troughs.
    iii. Ensure the tray is level to prevent uneven water heights between compartments, which can affect water exchange and animal conditions.
    iv. To check, place a bubble level across both the front–back and left–right axes of the tray, or visually compare water height across compartments after partial filling.
    v. If the tray is not level, adjust by repositioning risers or adding shims beneath the low corners until the water height is uniform across all compartments.
6. Install aeration.
  a. Place airstones in rear corner compartments (**Figure 1A**) to promote water mixing and oxygenation.

#### System validation before use

Before introducing animals:

1. Fill the tray with system water and allow it to run for at least 30 minutes.
2. Check all bulkheads and threaded connections for leaks.
3. Confirm equal water height across compartments.
  a. If water levels are unequal, verify that the tray is level, that standpipes are of equal height and fully seated, and that no obstructions are present in the outflow.
4. Confirm the bottom of the tray has not bowed and there are no gaps at the bottom of the acrylic of each compartment.
5. Measure temperature, salinity, pH, and dissolved oxygen to ensure consistency with home tanks.

#### CRITICAL

Confirm that there are still no gaps between the dividers, tray walls, and tray bottom now that the tray is bearing weight.

**Note:** Thread sealant tape and properly seated gaskets are typically sufficient to ensure watertight seals. If leaks are observed, drain the tray and apply a small amount of aquarium-safe silicone to the leaking interface and allow it to cure fully (≥48 hours) before refilling.

#### Usage notes

- Assign each compartment a grid coordinate (e.g. A1, B1) for ease of identification (**Figure 1A**).
- Transfer animals immediately after swabbing to their assigned compartment.
- Maintain animals in the tray only until genotyping results are available and new housing assignments are determined.

### Institutional permissions

The use of cephalopods in laboratory research is currently not regulated in the USA. However, Columbia University has established strict policies for the ethical use of cephalopods, including operational oversight from the Institutional Animal Care and Use Committee (IACUC). All of the cuttlefish used in this study were handled according to an approved IACUC protocol (AC-AABT8673). Experiments must be performed in accordance with local institutional and national guidelines and regulations.

### Key resources table

| REAGENT or RESOURCE | SOURCE | IDENTIFIER |
| --- | --- | --- |
| Chemicals, peptides, and recombinant proteins |  |  |
| APExBIO 2X SYBR Green qPCR Master Mix | APExBIO | K1070 |
| Agarose | Sigma-Aldrich | A7174-10G |
| 50x Tris-Acetate-EDTA (TAE) solution | Thermo Scientific | B49 |
| SYBR Safe DNA gel stain | Fisher Scientific | S33102 |
| 100 bp DNA ladder | New England Biolabs | N3231S |
| Nuclease-free water | Ambion | AM9930 |
| **Critical commercial assays** |  |  |
| Monarch Spin gDNA Extraction Kit | New England Biolabs | T3010L |
| **Experimental models: Organisms/strains** |  |  |
| *Ascarosepion bandense* | Columbia University | N/A |
| **Oligonucleotides** |  |  |
| See Table 1. |  |  |
| **Software and algorithms** |  |  |
| Geneious Prime 2025.2.2 | N/A | https://www.geneious.com |
| BLAST | N/A | https://blast.ncbi.nlm.nih.gov/Blast.cgi |
| QuantStudio Design & Analysis Software v2.2.0 | Applied Biosystems | N/A |
| IDT OligoAnalyzer | IDT | https://www.idtdna.com/calc/analyzer |
| **Other** |  |  |
| 20V Compact Drill | Dewalt | 885911723633 |
| 7/8" Hole Saw Bit | Grainger | 3ERF4 |
| Divider Box - 20" x 15" x 6" | ULINE | S-19496BLU |
| 1/2" FPT x FPT Bulkhead. Quantity: 2 | Lifegard Aquatics | R270900 |
| 1/2" Slip x 1/2" Male NPT Adapter. Quantity: 2 | Grainger | 22FJ19 |
| 1/2" x 10' PVC Schedule 40 Pipe<br>NOTE: there will be leftover PVC after construction | Grainger | 5AFJ2 |
| 1/2" FPT x 1/2" MPT 90° Elbow. Quantity: 2 | Grainger | 60UD05 |
| 1/2" FPT x 1/2" Slip 90° Elbow. Quantity: 2 | Grainger | 22FJ88 |
| Pipe Thread Sealant Tape (Teflon Tape) | Grainger | 3PDL5 |
| ¼" Clear Cast Acrylic Sheet (for lasercutting) - 32" x 18". Quantity: 2 | TAP Plastics | N/A |
| 1½" PVC Coupling (as risers). Quantity: 5 | Grainger | 22FM12 |
| ½" Slip x ½" Slip x ½" Slip Tee Pipe Fitting. Quantity: 2 | Grainger | 22FL02 |
| ½" Slip x ½" Slip 90° Elbow. Quantity: 2 | Grainger | 22FJ90 |
| ½" Threaded Pipe Nipple. Quantity: 4 | Grainger | 3TNV7 |
| Dow 795 Silicone Building Sealant - 10.3 Fluid Ounce Cartridge | PK Supplies | 795BL |
| Airstones | Bulk Reef Supply | 256276 |
| 1.5 ml Safe-Lock tubes | Eppendorf | 0030123611 |
| GeneMate SnapStrip 8-Strip Standard PCR Tubes with Individually Attached Flat Caps | GeneMate | 490003-706 |
| Nanodrop or equivalent | Thermofisher Scientific | ND-ONE-W |
| ThermoMixer or equivalent<br>Or a heat block + vortex | Eppendorf | 2231001127 |
| Mini Dry Bath or equivalent | Fisherbrand | 14-955-221 |
| Mini Dry Bath 1.5 ml Accessory Block | Fisherbrand | 14-955-224 |
| MicroAmp EnduraPlate Optical 384-Well Clear Reaction Plates or equivalent | Applied Biosystems | 4483285 |
| MicroAmp Optical Adhesive Film or equivalent | Applied Biosystems | 4311971 |
| QuantStudio 384-well qPCR machine or equivalent | Applied Biosystems | A28140 |
| Sartorius Picus 2 Electronic Multichannel Pipette (120 µL) or equivalent | Sartorius | LH-747421 |
| Sartorius Picus 2 Electronic Multichannel Pipette (10 µL) or equivalent | Sartorius | LH-747441 |
| Benchmark PLATEFUGE or equivalent | Benchmark Scientific | C2000 |
| 4" Deluxe Soft Rubber Brayer | Speedball | 4126 |
| Wide Mini-Sub Cell GT Cell DNA electrophoresis system or equivalent | Bio-Rad | 1704405 |

#### Step-by-step method details

##### Prepare tissue lysis buffer

*Timing: 15 min*

1. Label a series of 1.5 ml Safe-Lock tubes for the total number of animals being genotyped.
2. Calculate a master mix for the total number of samples (+10% for pipetting error):
3. Mix thoroughly by gentle pipetting or inversion.
4. Aliquot 200 µL of the master mix into each labeled tube and proceed directly to the next step.

| Reagent | 1X |
| --- | --- |
| Tissue lysis buffer<br>(from *Monarch Spin gDNA Extraction Kit*) | 200 μl |
| Proteinase K (20 mg/ml)<br>(from *Monarch Spin gDNA Extraction Kit*) | 3 μl |

**Note:** For DNA extractions from a piece of tissue (e.g. an arm tip) instead of a skin swab, use 10 μl of Proteinase K instead of 3 μl.

### Skin swabbing

*Timing: 1 hour (for 24 samples)*

In this step, skin cells are collected from live, unanesthetized cephalopods for DNA extraction. This step is best performed with two people: one to handle the animal and one to perform the swabbing.

1. Open a sterile foam swab and dip it into tank water to pre-soak.
2. Gently catch the animal using a soft net and hold it within the net (out of water), with its dorsal mantle facing up.
3. Using one side of the swab tip, gently swab the dorsal skin, moving systematically from left to right (**Figure 2A** and **Video 1**). **CRITICAL:** The dorsal skin is very delicate in young animals (<2 months old), so be extremely gentle. Avoid swabbing the skin all the way to the anterior end of the mantle, as this is the most delicate region of the skin. **Note**: In adults (>4 months), a mucus layer is present and additional pressure is required to collect sufficient cells.
4. Gently rotate the animal so its ventral side is facing upwards.
5. Rotate the swab 180° to use a clean side of the tip. Gently stroke the ventral surface in a left-to-right motion (**Figure 2B**). **Note:** Slightly greater pressure can be applied compared to the dorsal surface.
6. For swabbing hatchlings (**Figure 2C**):
  a. Place the hatchling on a microfiber towel suspended ∼5 mm below the surface of artificial seawater in a 5 L jug.
  b. Gently lift the towel edges to expose the mantle above water.
  c. Using a pre-soaked sterile foam swab, lightly swab the dorsal mantle, then rotate the animal and repeat on the ventral side with the opposite end of the swab.
  d. **CRITICAL**: Use extremely light pressure to avoid damaging the skin.
7. Place the swab face down in the corresponding Safe-Lock tube containing lysis buffer and leave it submerged (**Figure 2D**).
8. Return the animal to its designated tank or position in the housing tray (**Figures 2E** and **F**). **Note:** Recently swabbed skin may appear discolored or show transient markings. This is normal and typically resolves within minutes. Bright white patches indicate skin damage; in such cases, healing occurs within 24-48 hours, but reduced pressure should be used for subsequent animals.
9. Repeat steps 1-8 for each animal.

**Figure 2.**
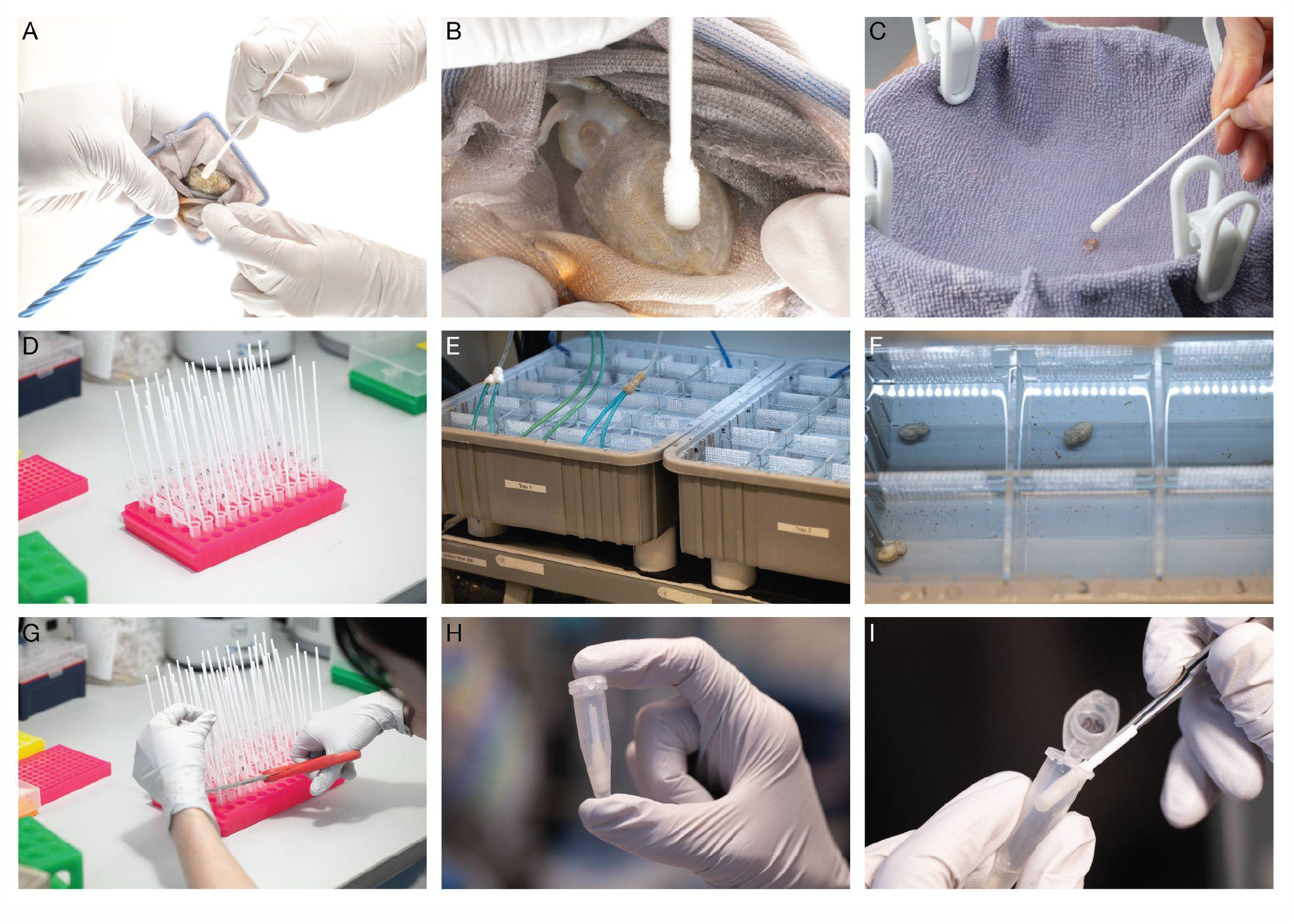
Skin swabbing and DNA extraction. A. Juvenile or adult cephalopods are caught in a net and gently swabbed on the dorsal mantle using a pre-soaked swab. B. The animal is rotated and the ventral surface is swabbed with slightly greater pressure. C. For hatchlings, animals are transferred onto a microfiber towel suspended in a jug of seawater such that the gills remain submerged. The dorsal and ventral surfaces are swabbed using extremely light pressure. D. Each swab is placed into a tube containing pre-mixed Tissue Lysis Buffer and Proteinase K. E. Immediately after swabbing, each animal is transferred to a housing tray (see “**Build a housing tray for temporary individual isolation**”). F. Animals are maintained in isolation to preserve identity. G. The swab shaft is cut using scissors. H. This allows the tube lid to close. I. After incubation with shaking for 1 hour, the swab is pressed against the side of the tube to recover residual liquid.

## DNA extraction

*Timing: 2 hours*

DNA is extracted from skin swabs collected in tissue lysis buffer.

1. Transfer the rack of Safe-Lock tubes (with foam swabs inside) to the bench.
2. For each tube, lift the swab a few millimeters and cut the shaft using clean scissors so that the tube lid can be fully closed (**Figures 2G and 2H**).
3. Incubate the tubes in a thermomixer at 56°C with shaking at 1,400 rpm for 1 hour. **Note:** If a thermomixer is not available, incubate tubes in a 56°C heat block and vortex vigorously every 5-10 minutes to facilitate lysis.
4. After incubation, briefly centrifuge the tubes to collect condensation and liquid at the bottom.
5. Using a gloved hand or forceps, grip the swab securely and press and drag it against the side of the tube to extract as much liquid as possible (**Figure 2I**).
6. Discard the swab and repeat for all samples.
7. Proceed with genomic DNA purification using the Monarch Spin gDNA Extraction Kit: **Note:** You can skip the RNase A step.
  a. Add 400 μl of gDNA Binding Buffer to each tube and mix thoroughly by pulse-vortexing for 5-10 seconds.
  b. Pour the lysate/binding buffer mix (∼600 μl) into a gDNA Purification Column pre-inserted into a collection tube.
  c. Centrifuge: first for 3 minutes at 1,000 x *g* to bind gDNA (no need to empty the collection tubes or remove from centrifuge) and then for 1 minute at maximum speed (> 12,000 x *g*) to clear the membrane. Discard the flow-through.
  d. Add 500 μl gDNA Wash Buffer, centrifuge immediately for 1 minute at maximum speed (12,000 x *g*), and discard the flow-through.
  e. Reinsert the column into the collection tube. Add 500 μl gDNA Wash Buffer and close the cap. Centrifuge immediately for 1 minute at maximum speed (>12,000 x *g*), then discard the collection tube and flow-through.
  f. Place the gDNA Purification Column in a DNase-free 1.5 ml microfuge tube.
  g. Add preheated (60°C) gDNA Elution Buffer directly to the center of the column membrane using the following volumes:
  h. Close the cap and incubate at room temperature for at least 5 minutes.
  i. Centrifuge for 1 minute at maximum speed (> 12,000 x *g*) to elute the gDNA.
8. **Note:** Typical yields from animals with a mantle length of 10-20 mm range from approximately 225 ng-1.1 µg total DNA. However, when eluted in 200 μl, the DNA concentration is usually below the accurate detection threshold of a Nanodrop. **CRITICAL:** There is no need to normalize the DNA concentrations.
9. For ease of pipetting, transfer samples into 8-strip PCR tubes (e.g. GeneMate SnapStrip 8-Strip Standard PCR Tubes with Individually Attached Flat Caps).

| Sample type | Elution volume |
| --- | --- |
| Hatchling | 50 μl |
| Juvenile | 200 μl |
| Adult | 50 μl |

### Pause point

DNA samples can be stored at −20°C if not used immediately.

### qPCR amplification of autosome and sex chromosome loci

*Timing: 3 hours*

Extracted genomic DNA is amplified via qPCR using primers targeting an autosomal locus, “A”, and a sex chromosomal locus, “S”, to obtain quantification cycle (Cq) values. Validated primer sets are provided in Table 1.

**Table 1.**
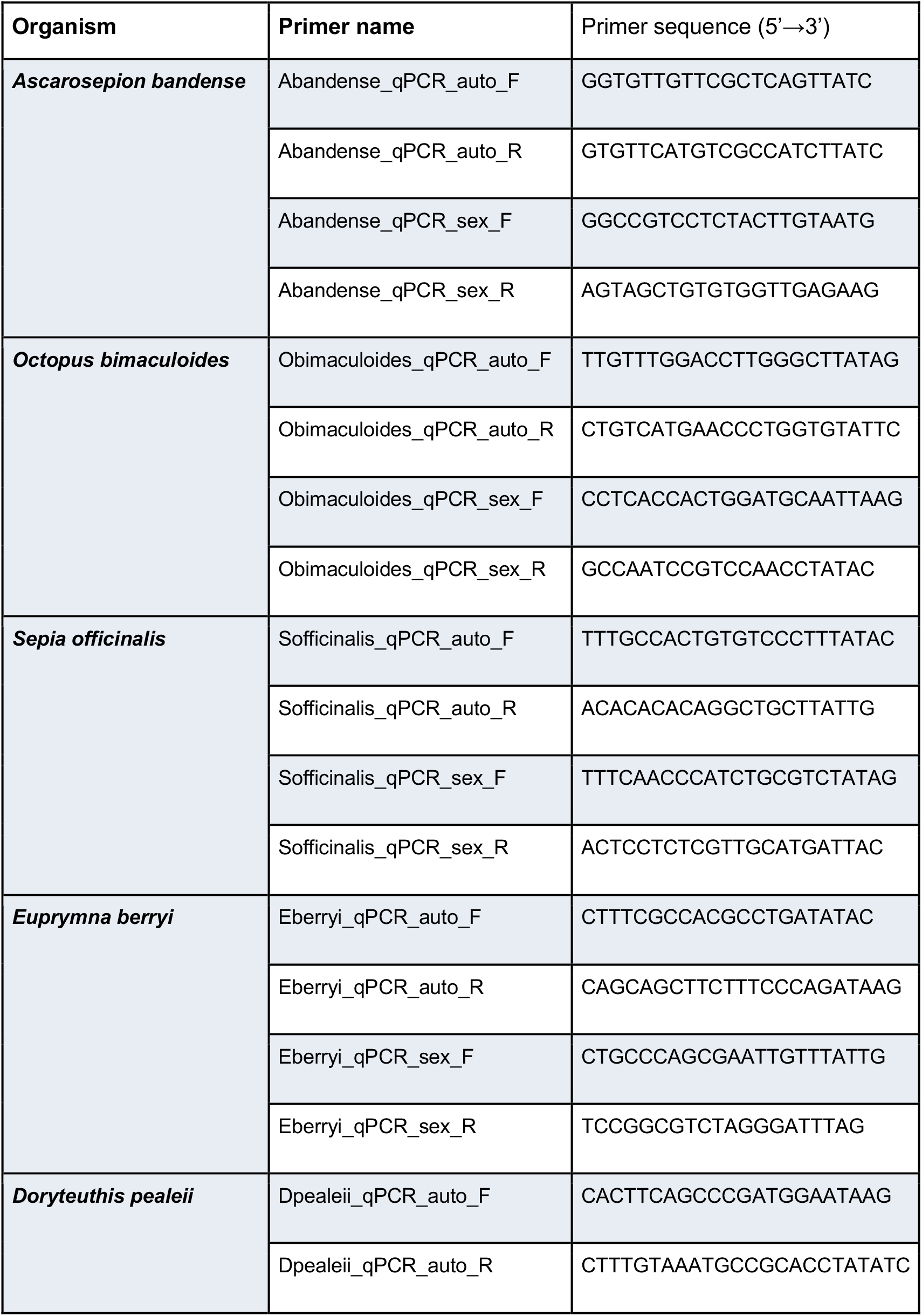
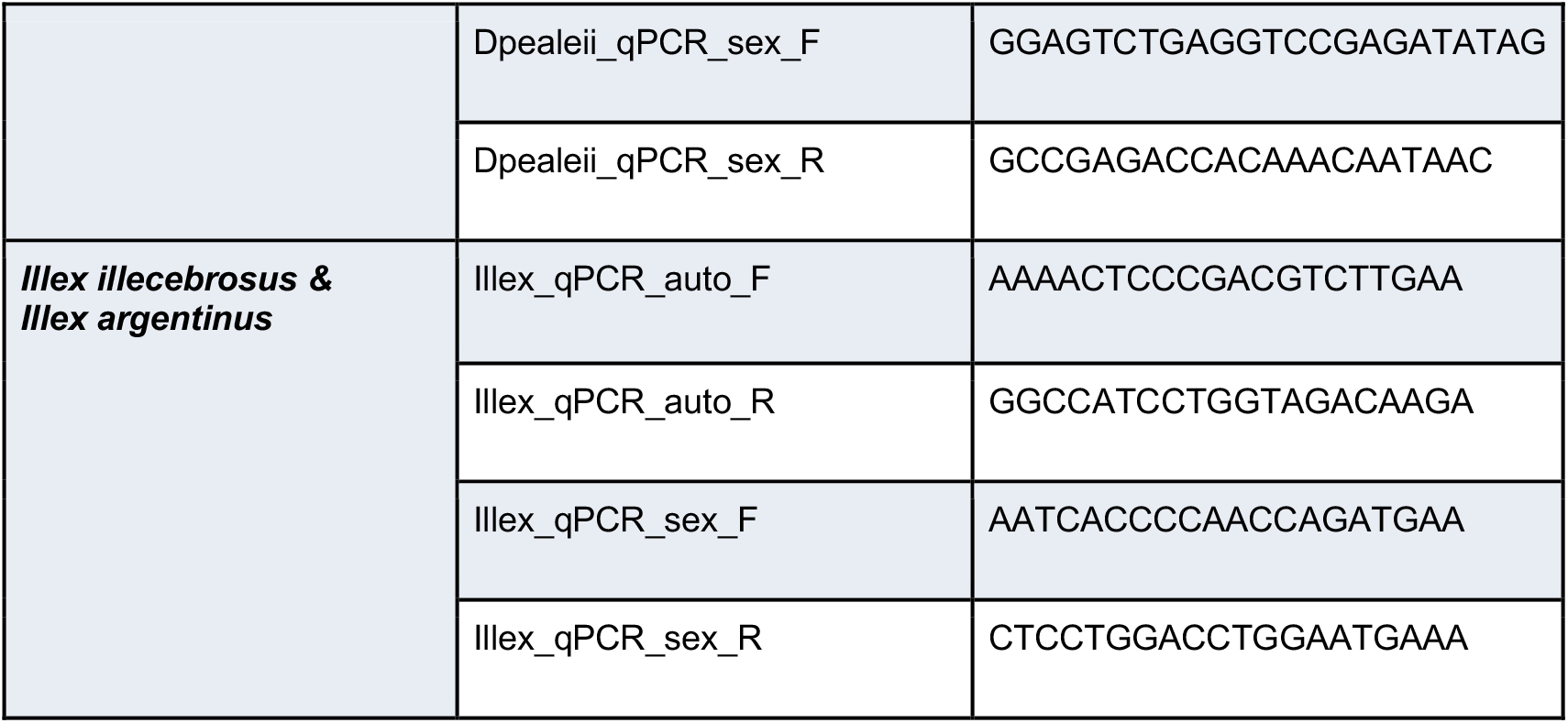
qPCR primer sequences for cuttlefish, octopus and squid species.

1. Prepare 20× primer mixes for autosomal (“A”) and sex chromosomal (“S”) loci in nuclease-free water. Each mix should contain forward and reverse primers at 5 µM each. Keep on ice. **Note:** Primer mixes can be stored at −20°C and reused for several months.
2. Thaw qPCR reagents on ice and prepare two master mixes “–A” & “S”. Reactions will have a final volume of 7 µL: 5 µL master mix + 2 µL sample DNA For N animals, prepare enough of **each** master mix (“A” & “S”) for 4×N reactions + 10% excess to account for pipetting loss. **Note:** For the most reliable qPCR results, work quickly and prepare master mixes immediately before adding sample DNA and running qPCR amplification. Minimize the time at room temperature by keeping reagents (and, optionally, the 384-well plate) on ice.
3. Using a multichannel pipette, load 5 µL master mix into wells of the 384-well plate. For each animal, perform 4 technical replicates per primer set. A 384-well plate layout can accommodate up to 48 animals (Figure 3). **Note**: An electronic repeating multichannel pipette is recommended to improve speed and dispensing accuracy.
4. Centrifuge the plate at 400 × g for 1 minute to collect the master mix at the bottom of the wells. **Note**: If an optically clear plate is used, visibly confirm the volume in each well is identical and no bubbles are trapped in the bottom of the well. Bubbles will interfere with the fluorescence readings of the qPCR machine.
5. Add 2 µL DNA sample to each well of the 384-well plate according to the schematic above. **Note**: Load the sample as a 2 µL bead along the sidewall of the tube using a repeating electronic multichannel pipette for the most consistent results. Continuously monitor the volume inside each pipette tip to ensure accurate dispensing. Examine pipette tips after each dispensed volume to ensure that sample fully entered the well and does not remain as a bead on the pipette tip. **Note**: Extra DNA samples can be stored at -20 °C for future use.
6. Gently tap the plate to bring droplets to the bottom. Seal with optical adhesive film, first by hand and then using a roller to ensure a tight seal.
7. Centrifuge the plate at 400 × g briefly to collect the volume at the bottom of the wells. To mix, hold the plate and use single swift motions to shake the contents toward the lid, and toward the bottom of the wells, alternately. Repeat 5 times.
8. Centrifuge the plate at 400 × g for 1 minute to collect the master mix at the bottom of the wells. **Note**: Confirm again that wells are free of bubbles and volumes are consistent.
9. Perform qPCR using an appropriate instrument with the following cycling conditions (**Figure 4**). **Note:** The following is for APExBIO SYBR green polymerase on QuantStudio 5, but follow your instrument and polymerase mix documentation if they differ:
  – **Initial denaturation:** 95°C for 2 min
  – **PCR** (45 cycles)
    – 95°C for 15 s
    – 60°C for 45 s
  – **Melt curve**
    – 95°C for 15 s
    – 60°C for 1 min
    – 95°C (at a rate of 0.06°C/s) for 15 s
    Set heated lid to 105°C and ramp rate to 1.6°C/s unless otherwise specified. ROX should be selected as the passive reference dye. **Note**: Regardless of manufacturer or model, ensure the instrument will calculate Cq values and record melt curve data.

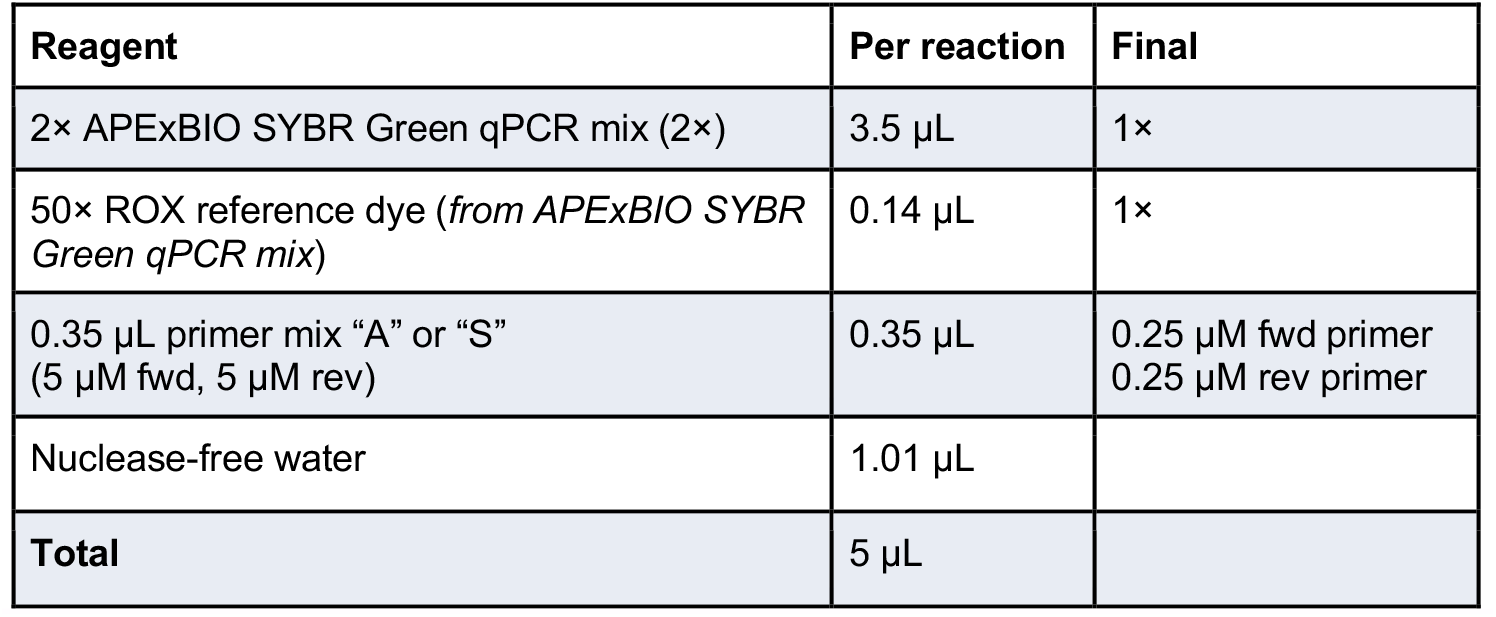

| Reagent | Per reaction | Final |
| --- | --- | --- |
| 2× APEXBio SYBR Green qPCR mix (2×) | 3.5 µL | 1× |
| 50× ROX reference dye (*from APEXBio SYBR Green qPCR mix*) | 0.14 µL | 1× |
| 0.35 µL primer mix “A” or “S”<br>(5 µM fwd, 5 µM rev) | 0.35 µL | 0.25 µM fwd primer<br>0.25 µM rev primer |
| Nuclease-free water | 1.01 µL |  |
| **Total** | 5 µL |  |

**Figure 3.**
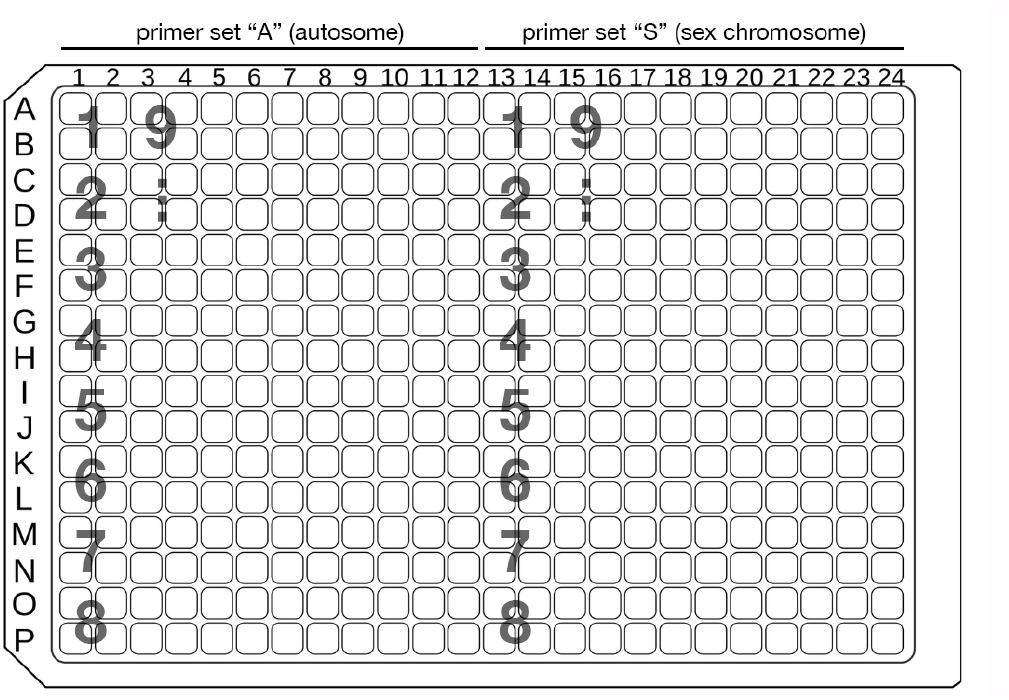
Recommended sample arrangement of a 384-well qPCR plate. The locations of samples for animals 1-9 are indicated by large gray letters. Four technical replicates are performed per primer set for each animal.

**Figure 4.**
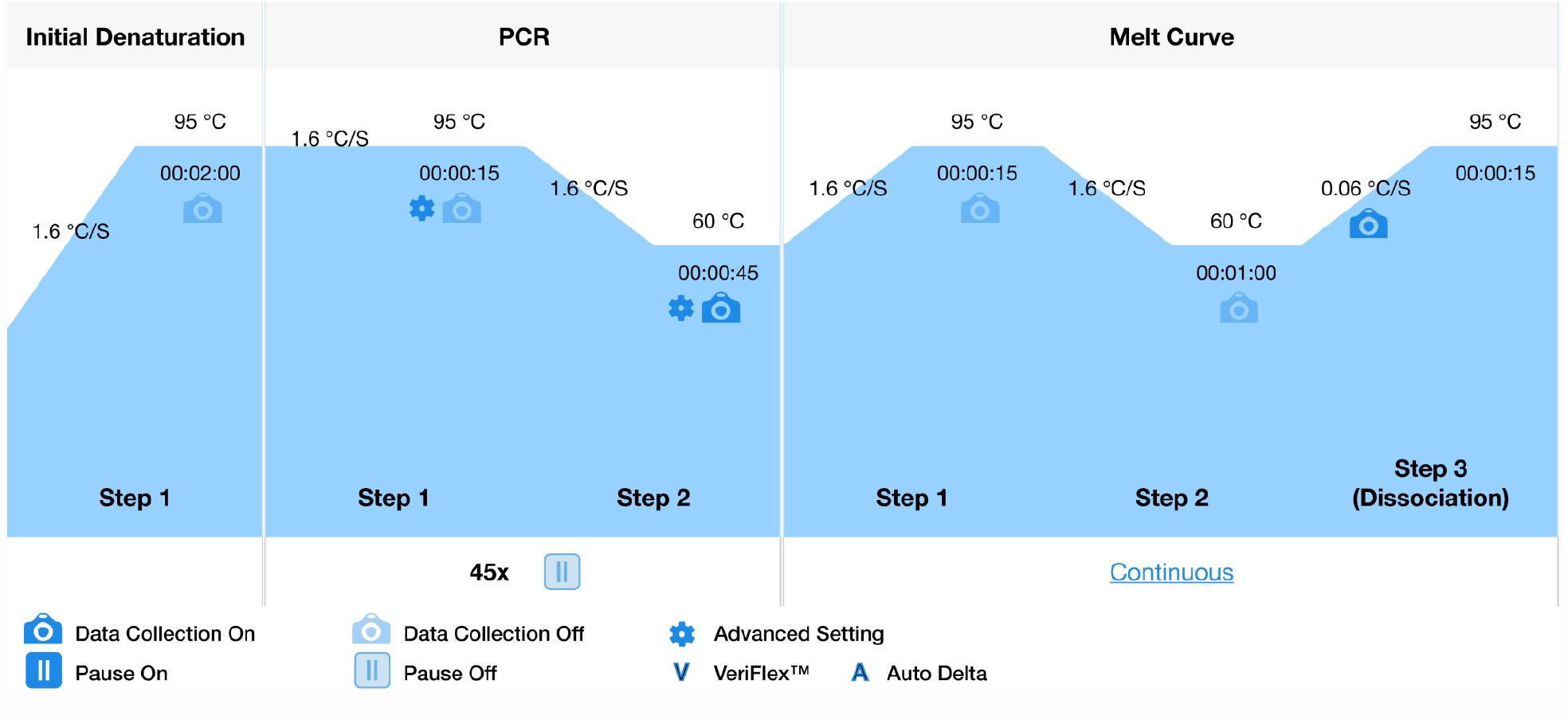
PCR protocol from QuantStudio 5 Design and Analysis software v2.2.0.

### qPCR analysis

*Timing: 30 min*

Analyze the qPCR results to determine which animals are males and females.

1. Inspect the melt curves of all samples to ensure a single melting point is observed for all samples of a given primer set (**Figure 5**). Discard any replicates with aberrant melt curves from the analysis. **Note**: For newly designed primers, confirm specificity by resolving amplicons on a 2% agarose gel and/or sequencing products (see “**Primer design**” and “**Primer validation**” sections). **Note**: Reasons for incorrect melting profile include low sample DNA concentration, or an off-target amplification. In the latter case, consider redesigning primers. Refer to “**Primer design**” and “**Primer validation**” sections.
2. Inspect the amplification plots of all samples to ensure a single slope giving a reliable Cq determination (**Figure 6**). Discard any replicates with aberrant amplification curves from the analysis. **Note:** Remove up to one outlier within a replicate set (e.g. Cq deviating by >1 cycle), which typically reflects a pipetting error. **Note**: Aberrant amplification curves may result from improperly loaded wells (e.g. bubbles), or from impure DNA preparation. For the latter, dilute the sample DNA 10-fold and repeat the amplification or prepare a new DNA sample. **Note**: Ensure all Cq values fall within the linear range of the primer pairs used. Otherwise, adjust template concentration accordingly. Refer to the “**Primer validation**” section. **Note**: For best results when genotyping multiple samples, use gDNA of roughly equal concentration. Preparing samples as suggested in the section “**DNA extraction**” should give reasonably similar concentrations for all sample types.
3. For each animal, calculate from the four replicates the mean Cq value and the standard error of the Cq value for each primer pair, A_Cq and S_Cq. **Note**: For *A. bandense*, following this protocol should produce Cq values in the range of 22-25 and standard error values below 0.1. The difference between a ZZ and Z0 individual is expected to be only 1 cycle; therefore, if standard error values rise above 0.25 cycles, the ability to discern males and females will diminish.
4. For each animal, calculate the Cq difference and the combined standard error:

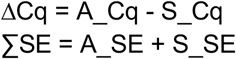

Normalize ΔCq values to be centered around zero, so that negative values predict females and positive values predict males. To estimate this offset, use the midpoint value between known males and known females. Alternatively, you may use the midpoint value between the lowest quartile (females) and highest quartile (males) ΔCq values.

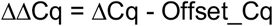

**Note:** Ideally Cq values including error bars should fall fully below or fully above zero and should form two clusters, male and female. Single outliers generally indicate a poor DNA sample. If data do not cluster into two groups, this may indicate insufficient pipetting accuracy when loading the master mix or DNA sample, or a sub-optimal primer set.
5. Assign sex based on ΔΔCq, as in **Figure 7**: Female: ΔΔCq < 0 Male: ΔΔCq > 0 Assign any individuals whose error bars cross the zero line as low confidence.

**Figure 5.**
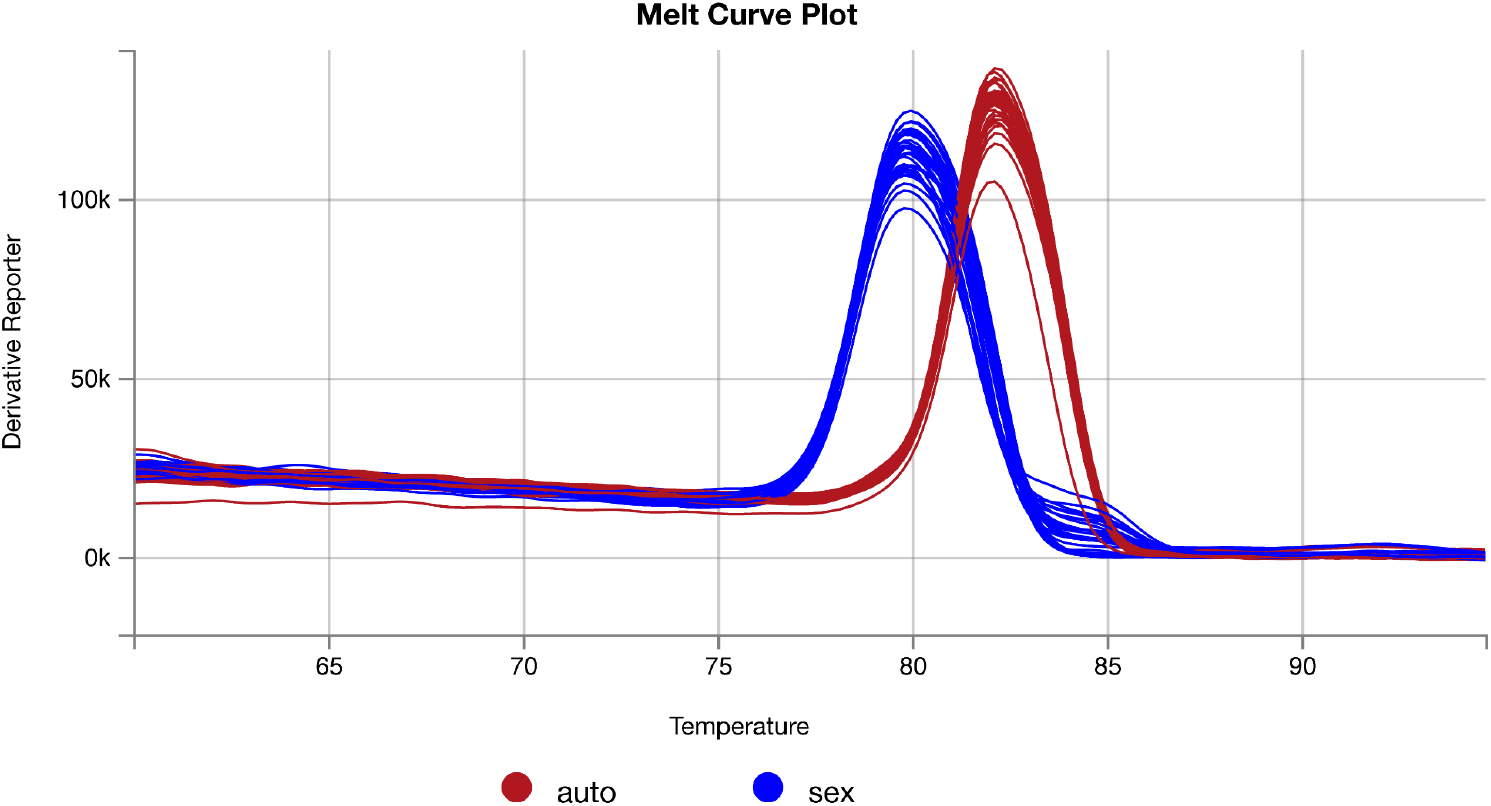
Melt curve plots of amplicons from autosome- and sex chromosome-targeting primers against 8 individuals.

**Figure 6.**
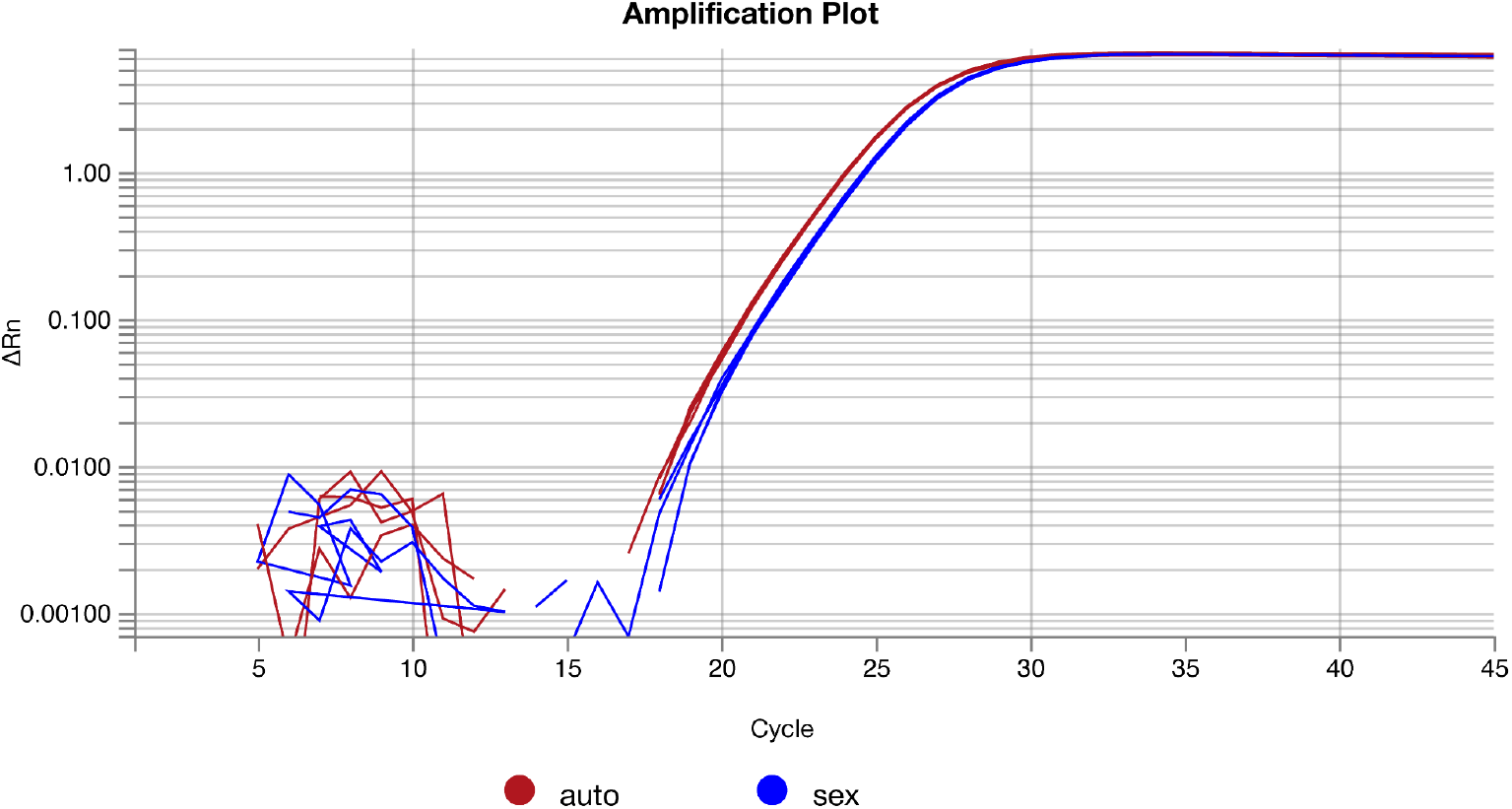
Amplification plots of four technical replicates from autosome- and sex chromosome-targeting primers for a single individual.

**Figure 7.**
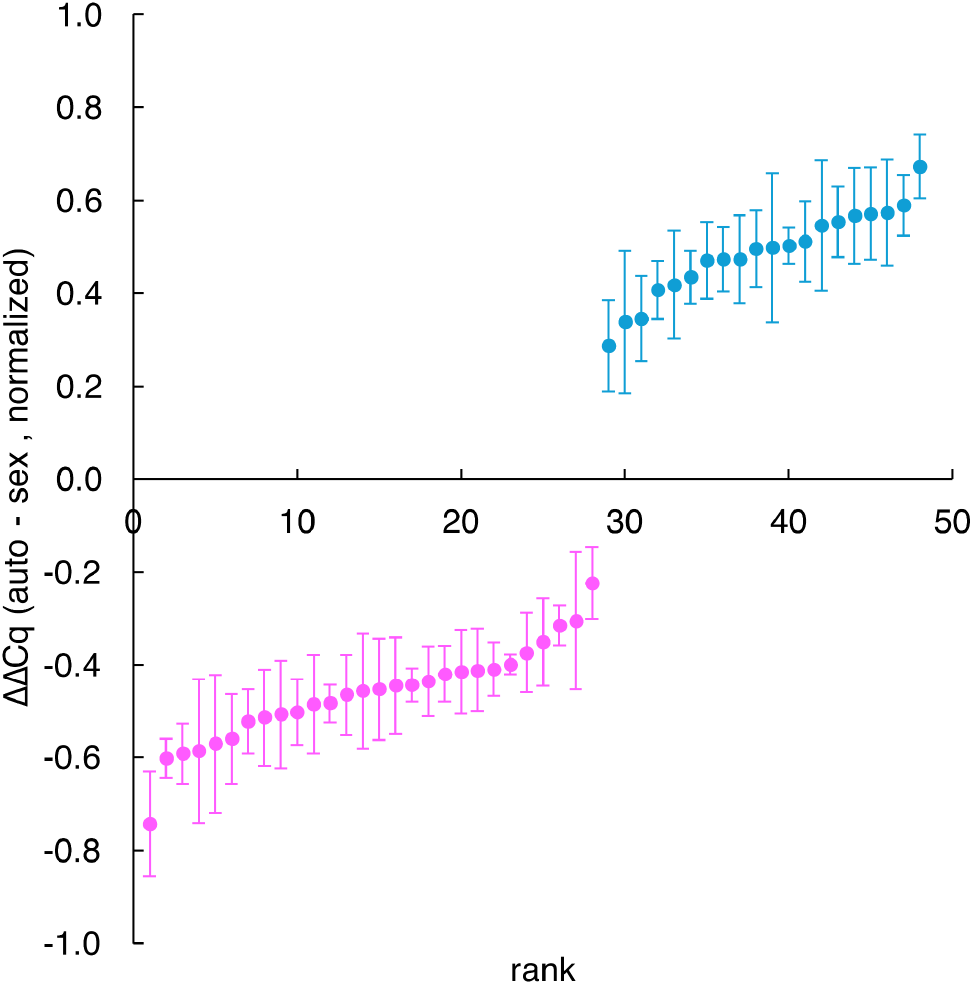
Normalized ΔΔCq values for 28 female (negative values, pink) and 20 male (positive values, blue) juvenile cuttlefish. Error bars represent standard error across 4 technical replicates.

### Expected outcomes

Following this protocol enables accurate prediction of sex in groups of cephalopods, such as the 48 dwarf cuttlefish shown here (**Figure 7**). We previously demonstrated that this approach achieves 100% accuracy, based on post-mortem validation of 81 dwarf cuttlefish^1^. In our hands, swabbing 48 animals, extracting genomic DNA, performing qPCR, and analyzing the results can all be completed within a single day (**Graphical Abstract**).

This protocol is broadly applicable across species and sample types. We provide validated primer sequences for seven cephalopod species: *O. bimaculoides, S. officinalis, A. bandense, E. berryi, D. pealeii, I. illecebrosus*, and *I. argentinus* (**Table 1**). For additional species, we outline a framework for designing and validating qPCR primers that detect the expected two-fold dosage difference between males and females (see “**Primer design and validation**”). For sample type, this protocol can be performed using non-invasive skin swabs from live animals (“**Skin swabbing**”) or using tissue samples such as arm tips or embryos. In all cases, genomic DNA is extracted using the Monarch Spin gDNA Extraction Kit and used as input for qPCR. Finally, this protocol can be used on living animals at a range of life stages, including hatchlings (**Figure 2C**), juveniles (**Figure 2A,B**), or adults, enabling accurate sex prediction well before sexual maturity and facilitating controlled housing and breeding.

### Limitations

This protocol is highly accurate when performed with validated primers, four technical replicates, and careful pipetting. Confidence in sex assignment is strengthened by analyzing multiple samples and observing clear separation between male and female ΔΔCq values. Accordingly, experiments should include at least four animals with both sexes represented.

This protocol requires access to a qPCR instrument. Laboratories without in-house access may use commercial qPCR services. However, cost and turnaround time may limit feasibility, particularly given the need to maintain animals in isolation following swabbing.

#### Troubleshooting

##### Problem 1: High variability in Cq values between technical replicates

**Potential solution:** Ensure Cq values fall within the validated linear range of the primers (see “**Primer validation**” section). Verify that pipettes are calibrated and pipetting technique is consistent. Confirm plates are centrifuged and reactions are thoroughly mixed prior to thermocycling. Minimize time at room temperature before qPCR.

##### Problem 2: High Cq values (late amplification)

**Potential solution:** Confirm samples fall within the linear range of each primer pair (see “**Primer validation**” section). If caused by low DNA concentration, re-swab animals using higher pressure to collect more cells (particularly in adults, which may have a mucus layer), elute DNA in a smaller volume, or concentrate existing samples (e.g. dry them in a speedvac and resuspend in a reduced volume).

##### Problem 3: ΔΔCq values do not separate into two clusters (male and female)

**Potential solution:** Confirm samples fall within the linear range of each primer pair (see “**Primer validation**” section). Normalize DNA sample concentrations so that the Cq values of all samples are similar (using the conversion that a two-fold dilution corresponds to ∼1 Cq shift). Note that minor deviations from 100% primer efficiency are tolerated when concentrations are similar, but larger differences in concentration can obscure clustering.

##### Problem 4: Amplification curves are preceded by a large peak and appear abnormal in slope or form

**Potential solution:** Some gDNA extractions may be contaminated with material from the animal that interferes with the qPCR reaction and is not removed by the purification kit. This can occur with degraded tissue, buccal samples, or overly concentrated swabs. Dilute samples in nuclease-free water and optionally heat at 98°C for 5 min before re-running. If issues persist, re-extract DNA or collect cleaner samples (e.g. gentle mantle swabs).

##### Problem 5: Incorrect sex prediction for an individual sample

##### Potential solution

If amplification and melt curves are normal, Cq values fall within the linear range, and technical replicates are tight (SEM < 0.1 cycles), consider biological explanations. These could include:

(1) copy number variation at the target locus; (2) discordance between genotype and phenotype (e.g. environmental factors altering sex determination, intersex animals, or behavioral sex mimicry); (3) an alternative sex determination system in the species. To distinguish these possibilities, repeat qPCR using independent primer pairs targeting different autosomal and sex-linked loci.

## Resource availability

### Lead contact

Further information and requests for any resources and reagents should be directed to and will be fulfilled by the lead contact, Tessa Montague.

### Technical contact

Questions regarding technical specifics of the protocol should be directed to the technical contact, Frederick Rubino.

## Materials availability

This study did not generate new unique reagents.

## Data and code availability

This study did not generate code. Raw data reported in this paper will be shared by the lead contact upon request.

## Acknowledgements

We would like to thank Richard Axel for valuable discussions, funding, and mentorship; Ruth Lehmann for funding and mentorship; Gabrielle Winters Bostwick and Sarah Detmering (Crook lab) for testing the protocol and providing feedback; and Thomas Barlow for photographing the swabbing process. F.A.R. is supported by the Helen Hay Whitney Foundation and was supported by the Whitehead Institute (through Dr. Ruth Lehmann); T.G.M. is supported by the HHMI Hanna H. Gray Fellowship; C.J.G., T.J.M., and T.G.M. were supported by the Howard Hughes Medical Institute (HHMI, through Dr. Richard Axel); and G.C.C., S.T.S., and A.D.K. were funded in part through NIH Awards R01HG010774 and R35GM148253.

## Author contributions

Conceptualization: F.A.R., C.J.G., G.C.C., A.D.K., T.G.M.; Methodology: F.A.R., C.J.G., T.J.M., T.G.M.; Investigation: F.A.R., C.J.G., T.G.M.; Formal analysis: F.A.R.; Resources: T.J.M., S.T.S.; Writing – original draft: F.A.R., T.J.M., T.G.M.; Writing – review & editing: F.A.R., C.J.G., T.G.M.; Visualization: F.A.R., T.J.M., T.G.M.; Project administration: A.D.K., T.G.M.; Supervision: A.D.K., T.G.M.

## Declaration of interests

The authors declare no competing interests.

## Declaration of generative AI and AI-assisted technologies in the manuscript preparation process

During the preparation of this work, the authors used ChatGPT for text editing suggestions to improve the clarity of the manuscript. After using this tool, the authors reviewed and edited the content as needed and take full responsibility for the content of the published article.

## Notes

### Competing Interest Statement

The authors have declared no competing interest.

