## Supplementary material for "Protocol for genotyping cephalopod sex using a skin swab and quantitative PCR": Table S1

| Chromosome | Orthogroup *O. bimaculoides* | Gene ID | Species | Sequence |
|---|---|---|---|---|
| chr1 (autosome) | OG0008421 | pred2_29441.1 | *Architeuthis dux* | MTKLEIITPFQLYFNPDLIFKKYHIWRLVTNFMYFGPIGFNFIFNIIFAYRYCRMLEEGSFRGRTSDFFFMFIFGGFVMIFITILIGNIAFLGNAFTIMLVYIWSRRNPYVRMNFFGLLNFQAPYLPWVLLAFSLLLGNSVAVDLMGILVGHLYYYLEDVFPHQVGGFKILRTPQFLKTLMDAAPEDPNYAPLPEDRPGGFNWGNQPQ |
| | | Dopeav2008472m.g | *Doryteuthis pealeii* | MAFQQEYMQMPPITRAYTTACVLTTIAVKLDIITPFQLYFNPDLIFKKYHIWRLVTNFMYFGPIGFNFIFNIIFAYRYCRMLEEGSFRGRTSDFFFMFIFGGFIMILITILIGNIAFLGNAFTIMLVYIWSRRNPYVRMNFFGLLNFQAPYLPWVLLAFSLLLGNSVAVDLMGILVGHLYYYLEDVFPHQVGGFKILRTPQFLKMLMDAAPEDPNYQPLPEDRPGGFNWGNQPQ* |
| | | g1053.t1 | *Euprymna scolopes* | MAFQQEYMQMPPITRAYTTACVLTTIAVKLDIITPFQLYFNPDLIFKKYHVWRLVTNFMYFGPIGFNFIFNIIFAYRYCRMLEEGSFRGRTSDFFFMFIFGGFIMIIITILIGNIAFLGNAFTIMLVYIWSRRNPYVRMNFFGLLNFQAPYLPWVLLAFSLLLGNSVAVDLMGILVGHLYYYLEDVFPHQVGGFKILRTPQFLKTLMDAAPEDPNYQPLPEDRPGGFNWGNQGQ |
| | | LOC_00005141-mRNA-1 | *Illex illecebrosus* | MAFQQEYMQMPPITRAYTTACVLTTIAVKLEIITPFQLYFNPDLIFKKYHIWRLVTNFMYFGPIGFNFIFNIIFAYRYCRMLEEGSFRGRTSDFFFMFIFGGFIMILITILIGNIAFLGNAFTIMLVYIWSRRNPYVRMNFFGLLNFQAPYLPWVLLAFSLLLGNSVAVDLMGILVGHLYYYLEDVFPHQVGGFKILRTPQFLKTIMDAAPEDPNYAPLPEDRPGGFNWGNQPQ |
| | | obimac_0000979.2 | *Octopus bimaculoides* | MAAQTFQQEYMQMPPITRAYTTACVLTTIAVKLDIITPLQLYFNPDLIFKKYQIWRLLTNFMYFGPIGFNFIFNIIFAYRYCRMLEEGSFRGRTSDFFFMFLFGGFIMIINTILIGNIAFLGNAFTIMLVYIWSRRNPYVRMNFFGLLNFQAPYLPWVLLTFSLLLGNSVAVDLMGILVGHLYFYLEDVFPLQMGGFKILRTPRFLKYLMDTTVEDPNYQPFPEDRPGGFDWGNNGN |
| | | XP_029634577.2 | *Octopus sinensis* | MRTCQALEARMRINTWHSAMAAQTFQQEYMQMPPITRAYTTACVLTTIAVKLDIITPLQLYFNPDLIFKKYQIWRLLTNFMYFGPIGFNFIFNIIFAYRYCRMLEEGSFRGRTSDFFFMFLFGGFIMIINTILIGNIAFLGNAFTIMLVYIWSRRNPYVRMNFFGLLNFQAPYLPWVLLTFSLLLGNSVAVDLMGILVGHLYFYLEDVFPLQMGGFKILRTPRFLKYLMDTTVEDPNYQPFPEDRPGGFDWGNNGN |
| | | rna-SPHA_31531 | *Sepia esculenta* | MPPITRAYTTACVLTTIAVKLDIITPFQLYFNPDLIFKKYHIWRLVTNFMYFGPIGFNFIFNIIFAYRYCRMLEEGSFRGRTSDFFFMFLFGGFIMILITILIGNIAFLGNAFTIMLVYIWSRRNPYVRMNFFGLLNFQAPYLPWVLLAFSLLLGNSVAVDLMGILVGHLYYYLEDVFPHQVGGFKILRTPQFLKTIMDAAPEDPNYQPLPEDRPGGFNWGNQPQ* |
| | | rna-SPHA_31531 | *Sepia officinalis* | MPPITRAYTTACVLTTIAVKLDIITPFQLYFNPDLIFKKYHIWRLVTNFMYFGPIGFNFIFNIIFAYRYCRMLEEGSFRGRTSDFFFMFLFGGFIMILITILIGNIAFLGNAFTIMLVYIWSRRNPYVRMNFFGLLNFQAPYLPWVLLAFSLLLGNSVAVDLM |
| chr1 (autosome) | OG0009041 | pred2_18336.1 | *Architeuthis dux* | MPHIDNDIKLDFKDVLLRPKRSTIKSRADVDLGREFIFRNSGKTYQGIPIMASNMDTVGTFEMANELSKHRVFTVIHKHYSLDAWKEFAANNPDTLKNIAVSSGMAGGDREKLEAIVNAIPGLHYICLDVANGYSEYFVQFVKDVRKKFPHHVILAGNVVTGEMVEELILSGADIIKVGIGPGSVCTTRKKTGIGYPQLSAVIECADAAHGLGGHIISDGGCTCPGDVAKAFGAGADFVMVGGMLAGHTESGGEMIEKNGKKFKLFYGMSSATAMQKHVGKVAEYRASEGKTVEIAYRGDVQVTMLDILGGIRSTCTYVGASKLKELSRRTTFIRVTQQVNEVFTPFTTAP |
| | | Dopeav2098899m.g | *Doryteuthis pealeii* | MPHIDNDIKLDFKDVLLRPKRSTIKSRADVDLGREFIFRNSGKTYKGIPIMASNMDTVGTFEMANQLAKHGVFTVIHKHYSLDAWKEFAAKNPETLNNIAVSSGMAAGDFEKLEAVINSIPALQYICLDVANGYSEYFVQFVRDVRKRFPKHIILAGNVVTGEMVEELILSGADIIKVGIGPGSVCTTRKKTGIGYPQLSAVIECADAAHGLGGHIISDGGCTCPGDVAKAFGAGADFVMIGGMLAGHTESGGEMIEKNGKKFKLFYGMSSATAMQKHSGKVAEYRASEGKTVEIEYRGDVQGTILDVLGGLRSTCTYVGASKLKELSRRTTFIRVTQQVNEVFSPFTTAH |
| | | g14646.t1 | *Euprymna scolopes* | MPHIDNDIKLDFKDVLLRPKRSTIKSRADVDLVREFIFRNSGRTYQGIPVMASNMDTVGTFEMAKQLSKHRVFTVIHKHYSLEAWKEFAAKNPETLKNTAVSSGMAAGDLEKLAAIVNAIPDLHYICLDVANGYSEYFVQFVRDVRKKFPNHVIMAGNVVTGEMVEELILSGADIIKVGIGPGSVCTTRKKTGIGYPQLSAVIECADAAHGLGGHIISDGGCTCPGDVAKAFGAGADFVMVGGMLAGHTESGGDIVEKNGKKFKLFYGMSSATAMQKHSGKVAEYRASEGKTVEIEYRGDVQGTMLDVLGGLRSTCTYVGASKLKELSRRTTFIRVTQQVNEVFSPFTKAP |
| | | LOC_00006323-mRNA-1 | *Illex illecebrosus* | MPRIDNDIKLDFKDVLLRPKRSTIKSRADVDLGREFIFRNSGKTYEGIPIMASNMDTVGTFEMANELAKHRVFTVIHKHYSLDAWKEFAANNPDTVKNIAVSSGMAGGDLEKLEAVVNAIPDLNYICLDVANGYSEFFVQFVRDVRKKFPHHVILAGNVVTGEMVEELILSGADIIKVGIGPGSVCTTRKKTGIGYPQLSAVIECADAAHGLGGHIISDGGCTCPGDVAKAFGAGADFVMVGGMLAGHTESGGEMIEKNGKKFKLFYGMSSATAMHKHSGKVAEYRASEGKTVEIAYRGDVQATILDVLGGIRSTCTYVGASKLKELSRRTTFIRVTQQVNEVFSPFTTAP |
| | | obimac_0001039.2 | *Octopus bimaculoides* | MPRIDSDIKLDFKDVLLRPKRSTIKSRADVDLTREFIFRNSGQTYNGIPVMASNMDTVGTFEMAITLAKYGLFTTIHKHYSIEAWKEFAANHPDQLSNIAASSGMAAGDLQKLESIIEAVPALRYICLDVANGYSEYFVQFLRDVRKKFPSHVIMAGNVVTGEMVEELILSGADIIKVGIGPGSVCTTRKKTGIGYPQLSAVIECADAAHGLGGHIISDGGCTCPGDVAKAFGAGADFVMIGGMLAGHTESGGEMIEKNGKKFKLFYGMSSATAMQKHAGHVAEYRASEGKTVEIEYRGDASHTVQDILGGIRSTCTYVGASKLKELSRRTTFIRVTQQMNEVFSPFNKSC |
| | | XP_029641704.2 | *Octopus sinensis* | MRPEVLGEWVDLTREFIFRNSGQTYNGVPVMASNMDTVGTFEMAITLAKYGLFTTIHKHYSIDAWKEFAAKHPDKLNNIAASSGMAAGDLQKLESIIEAVPDLRYICLDVANGYSEFFVQFLRDVRKKFPSHVIMAGNVVTGEMVEELILSGADIIKVGIGPGSVCTTRKKTGIGYPQLSAVIECADAAHGLGGHIISDGGCTCPGDVAKAFGAGADFVMIGGMLAGHTESGGEMIEKNGKKFKLFYGMSSATAMQKHAGHVAEYRASEGKTVEIEYRGDAAHTVQDILGGIRSTCTYVGASKLKELSRRTTFIRVTQQMNEVFSPFNKSS |
| | | rna-SPHA_21857 | *Sepia esculenta* | VDLVREFIFRNSGQTYRGIPIMASNMDTVGTFEMAIQLSKHGVFTVIHKHYSLEAWKEFATKNPETVNNIAVSSGMAGGDLEKLESIVNAIPDLHYICLDVANGYSEYFVQFVRDVRKKFPKHVIMAGNVVTGEMVEELILSGADIIKVGIGPGSVCTTRKKTGIGYPQLSAVIECADAAHGLGGHIISDGGCTCPGDVAKAFGAGADFVMVGGMLAGHTESGGEMIERNGKKFKLFYGMSSATAMKKHAGKVAEYRASEGKSVEIEYRGDVQGTILDILGGIRSTCTYVGASKLKELSRRTTFIRVTQQLNEVFSPFTKAP |
| | | rna-SPHA_21857 | *Sepia officinalis* | MPHIDNDIKLDFKDVLLRPKRSTIKSRADVDLVREFLFRNSGQTYRGIPIMASNMDTVGTFEMAIQLSKHGVFTTIHKHYSLEAWKEFATKNPETVNAGNVVTGEMVEELILSGADIIKVGIGPGSVCTTRKKTGIGYPQLSAVIECADAAHGLGGHIISDGGCTCPGDVAKAFGAGADFVMVGGMLAGHTESGGEMIERNGKKFKLFYGMSSATAMKKHAGKVAEYRASEGKSVEIEYRGDVQGTILDVLGGIRSTCTYVGASKLKELSRRTTFIRVTQQLNEVFSPFTTAP |

| | | | | |
|---|---|---|---|---|
| chr1 (autosome) | OG0009283 | pred2_28664.1 | *Architeuthis dux* | MSYGRRGPSELDLEGMVSLKVDNLTYRTTPEDLRRAFEKYGDVGDVYIPRDRFTRESRGFAFVRYHDKRDADDAMDAMDGAIMDGRELRVQMARYGRPADPYRRGPPPRRFGGGGGGGGGGGGGGGGGGGGGGYSSYRSRRRSYSRSRSRSRSRSSRRRRRSYSRSRSRSRSRSYKSKSRSRSRSPSYSRSSRSKHYSRSPTPPSKGSGSKKRSSGGSRRSTRSRSPSRDDRKNSRSRSRSPSPARSSEGGRGESQGRGSGGPTGGGGASAGASGSGSSLAPAASTAGGGAGGAPRENSSKSRSRSRSRSRSRSRSRSRSNSRSPRYASVSPEGSPEANSNNHKDQQQADEED |
| | | Dopeav2005453m.g | *Doryteuthis pealeii* | MSYGRRGPSELDLEGMVSLKVDNLTYRTTPEDLRRAFEKYGDVGDVYIPRDRFTRESRGFAFVRYHDKRDADDAMDAMDGAIMDGRELRVQMARYGRPADPYRRGPPPRRFGGGGGGGGGYSSYRSRRRSYSRSRSRSRSRSSRRRRRSYSRSRSRSRSRSYKSRSRSRSPSYSRSSRSKHYSRSPTPPSKGSGSKKRSSGGSRRSTRSRSPSRDDRKSRSRSRSPSPARSSEGGRGGSQGRSSGPAGGVGSAGGSGSSGLAPAAGTASGGPPRENSKSRSRSRSRSRSRSRSRSRSRSHSRSRSRSPRSRSCSPEGSPDANSNNHKDKQQVDEED |
| | | g752.t1 | *Euprymna scolopes* | MSYGRRGPSELDLEGMVSLKVDNLTYRTTPEDLRRAFEKYGDVGDVYIPRDRFTRESRGFAFVRYHDKRDADDAMDAMDGAIMDGRELRVQMARYGRPADPYRRGPPPRRFGGGGGGGGGGGGGGGYSSYRSRRRSYSRSRSRSRSRSSRRRRRSYTRSRSRSRSRSYKSRSRSRSPSYSRSSRSKHYSRSPTPPSKGSGSKKRSSGGSRRSTRSRSPSRDDRKSRSRSRSASPARSSEGGRGGSQTRGSSGPPASSGATASASGNLTPSGSATASGGAPRENSSKSRSRSRSRSRSRSRSRSPHSRSRSRSRSRSCSPQGSPDANSNNHKDKQEVDEED |
| | | LOC_00005496-mRNA-1 | *Illex illecebrosus* | MSYGRRGPSELDLEGMVSLKVDNLTYRTTPEDLRRAFEKYGDVGDVYIPRDRFTRESRGFAFVRYHDKRDADDAMDAMDGAIMDGRELRVQMARYGRPADPYRRGPPPRRFGGGGGGGGGGGGGGGGGGGGYSSYRSRRRSYSRSRSRSRSRSSRRRRRSYTRSRSRSRSRSYKSRSRSRSPSYSRSSRSKHYSRSPTPPSKGSGSKKRSSGGSRRSTRSRSPSREERKTSRSRSRSPSPARSSEGGRAGSQGRGSSGPAAGAGAGGSAGASGSSSSLVPAAGTTSGGAGAARENSSKSRSRSRSRSRSRSRSRSRSRSRSHSRSPRSRSVSPEGSPEANSNNHKDQQQADEED |
| | | obimac_0002028.1 | *Octopus bimaculoides* | MSYVRRGPSELDLEGMVSLKVDNLTYRTTPEDLRRAFEKYGDVGDVYIPRDRFTRESRGFAFVRYHDKRDADDAMDAMDGAMMDGRELRVQMARYGRPADPYRRGPPPRRFGGGGYSSYRSRRRSYTRSRSRSRSRSSRRRRRSYSRSRSRSRSRSYKSRSRSRSASYSRSSRSKHYSRSPSPSSKGKKRSSASSRRSTRSRSASRERKSHSRSRSPVSPRRSPDRCSEGRDGSESGAGANAGGGGGGGSSGSVPRENSKSRSRSRTPHSRSCSPEALSDRGSNDHNDLEQQVEEED |
| | | XP_029633039.1 | *Octopus sinensis* | MSYVRRGPSELDLEGMVSLKVDNLTYRTTPEDLRRAFEKYGDVGDVYIPRDRFTRESRGFAFVRYHDKRDADDAMDAMDGAMMDGRELRVQMARYGRPADPYRRGPPPRRFGGGGYSSYRSRRRSYTRSRSRSRSRSSRRRRRSYSRSRSRSRSRSYKSRSRSRSASYSRSSRSKHYSRSPSPSSKGKKRSSASSRRSTRSRSASRERKSHSRSRSPVSPRRSPDRCSEGRDGSESGAGANAGGGGGGGSSGSVPRENSKSRSRSRTPHSRSCSPEALSDRGSNDHNDLEQQVEEED |
| | | rna-SPHA_39875 | *Sepia esculenta* | MSYGRRGPSELDLEGMVSLKVDNLTYRTTPEDLRRAFEKYGDVGDVYIPRDRFTRESRGFAFVRYHDKRDADDAMDAMDGAVMDGRELRVQMARYGRPADPYRRGPPPRRFGGGGGGGGGGGYSSYRSRRRSYSRSRSRSRSRSSRRRRRSYTRSRSRSRSRSYKSRSRSRSPSYSRSSRSKHYSRSPTPPSKGSGSKKRSSGGSRRSTRSRSPSRDDRKSRSRSRSPGSPARSSEGGRGGSQGRSSGPAGGGGSTGGSGSGTLAPAAGTASGGPPRENSSKSRSRSRSRSRSRSRSRSRSRSRSRSPRSRSCSPEGSPDANSNNHKDKQQIDEED |
| | | rna-SPHA_39875 | *Sepia officinalis* | MSYGRRGPSELDLEGMVSLKVDNLTYRTTPEDLRRAFEKYGDVGDVYIPRDRFTRESRGFAFVRYHDKRDADDAMDAMDGAVMDGRELRVQMARYGRPADPYRRGPPPRRFGGGGGGGGGGGGGYSSYRSRRRSYSRSRSRSRSRSSRRRRRSYTRSRSRSRSRSYKSRSRSRSPSYSRSSRSKHYSRSPTPPSKGSGSKKRSSGGSRRSTRSRSPSRDDRKSRSRSRSPGSPARSSEGGRGGSQGRSSGPVGGGSSAGGSGSGTLAPAAGTASGGAPRENSSKSRSRSRSRSRSRSRSRSRSRSRSRSPRSRSCSPEGSPDANSNNHKDKQQIDEED |
| chr1 (autosome) | OG0009320 | pred2_29997.1 | *Architeuthis dux* | MLPGVGVFGTTATIQSFVPILKVCGFPVVALWGSENLDAQELATQLDIPFSTRKVDDVLLRKDVDLVVIGCPPHSQWFIAVKALGIGKHVLCSAPSGPMQLNAQHMVKAARYYPRLMSLMCYGLRFLPTIVKMKKMIEDGCLGNITICEVKVHYGGLPKEKYDWMCDEGMGGGVLNTIGSNIIDVITFLTKERAVRVHGMLKTYTKQTQNIKGIREITSDDFCTFQMELDKGACVTVTLNSHIPGQFVQEIVICGQKGRLIARGSDLYEQRLNATRESLIHFDPIKEEERYGISPKARTEIPSPYLKGLIHMIEAVKDAFEKEEERQNWQAEPVASASNFEDALYVQTVVDAIRKSNKTKQWTKVDVSVEEPEPNSNNMLSDQMRRSTFSLQ |
| | | Dopeav2006705m.g | *Doryteuthis pealeii* | MLPGVGVFGTTATIQSFVPILKVCGFPVVALWGRENLDAQELATQLDIPFSTRKVDDVLLRKDVDLVVIGCPPHSQCFIAVKALGIGKHVLCSAPSGPMQLNVQRMVKAARYYPRLMSLMCYGLRFLPTIVKMKKMIEEGCLGSITICEVKVHYGGLPKEKYDWMCDEGMGGGVLNTIGSNIIDVITFLTKERAVRVHGMLKTYTKQTQNIKGIREITSDDFCTFQMELDKGACVTVTLNSHIPGEFVQEIVICGQKGRLIARGSDLYEQRLNATRESLIHFDAIKEEEKYGISPKARTEIPIPYLKGLIHMIEAVKDAFEKEEERQNWQGEPVASASNFEDALYVQTVVDAIRKSNKTKQWTKVDVSVEEPEPNSNNMLSDQMRRSTFSLQ |
| | | g828.t1 | *Euprymna scolopes* | MLPGVGVFGTTATIQSFVPILKVCGFPVVALWGSENLDAQELATQLDIPFSTRKVDDVLLRKDVDLVVIGCPPHSQWFIAVKALGIGKHVLCSAPSGPMQLNAQHMVKAARYYPRLMSLMCYGLRFLPTIIKMKTMIEEGCLGSITICEVKVHYGGLPKEKYDWMCDEGMGGGVLNTIGSNIIDVITFLTKERALRVHGMLKTYTKQTQNIKGIREITSDDFCTFQMELDKGACVTVTLNSHIPGQFVQEIVICGQKGRLIARGSDLYEQRLNATRESLIHFDPIKEEERYGISPKARTEIPSPYLKGLIHMIEAVKDAFEKEEERQNWQAEPVASASNFEDALYVQTVVDAIRKSNKTKQWTKVDVSVEEPEPNSNNMLSDQMRRSTFSLQ |
| | | LOC_00005427-mRNA-1 | *Illex illecebrosus* | MLPGVGVFGTTATIQSFVPILKVCGFPVVALWGSENLDAQELATQLDIPFSTRKVDDVLLRKDVDLVVIGCPPHSQWFIAVKALGIGKHVLCSAPSGPMQLNAQHMVKAARYYPRLMSLMCYGLRFLPTIIKMKKMIEEGCLGSITICEVKVHYGGLPKEKYDWMCDEGMGGGVLNTIGSNIIDVITFLTKERAVRVHGMLKTYTKQTQNIKGIREITSDDFCTFQMELDKGACVTVTLNSHIPGQFVQEIVICGQKGRLIARGSDLYEQRLNATRESLIHFDPIKEEERYGISPKARTEIPSPYLKGLIHMIEAVKDAFEKEEERQNWQTEPVASASNFEDALYVQTVVDAIRKSNKTKQWTKVDVSVEETNSNNMLSDQMRRSTFSLQ |
| | | obimac_0001416.1 | *Octopus bimaculoides* | MLPGVGVFGTTATIQSFVPILKVCGFPVVALWGSENLDARTLAAELDIPFSTHKVDDVLLRKDVDLVVIGCPPHSQSYIAVKALGIGKHVLCSSPSGPMQLNALLMVKAARYYPRLMSLMCYGLRFLPTIVKMKKIIEDGCLGNITICEVKVHYGGLPKEKYDWMCDEGMGGGVLNTFGSNIIDVITFLTSERAVRVHGMLKTYTKQTQNIKGIREITSDDFCTFQMELDKGACVNVTLNSHITGQFVQEILVCGRKGRLIARGSDLYEQRLNATRESLIHFDPIKEEERYGISPKARTEIPSPYLKGLIHMIEAVKDAFEKEEERQNWHPEPVASASNFEDALYVQAVVDAIRKSNKTKQWTKVDVSVNEPEPEPNSNSMSLDYVRRSTIFLH |

| | | | | |
|---|---|---|---|---|
| | | XP_029644474.1 | *Octopus sinensis* | MLPGVGVFGTTATIQSFVPILKVCGFPVVALWGSENLDARTLAAELDIPFSTHKVDDVLLRKDVDLVVIGCPPHSQSYIAVKALGIGKHVLCSSPSGPMQLNALLMVKAARYYPRLMSLMCYGLRFLPTIVKMKKIIEEGCLGNITICEVKVHYGGLPKEKYDWMCDEGMGGGVLNTFGSNIIDVITFLTSERAVRVHGMLKTYTKQTQNIKGIREITSDDFCTFQMELDKGACVNVTLNSHITGQFVQEILVCGRKGRLIARGSDLYEQRLNATRESLIHFDPIKEEERYGISPKARTEIPSPYLKGLIHMIEAVKDAFEKEEERQNWHPEPVASASNFEDALYVQAVVDAIRKSNKTKQWTKVDVSVNEPEPEPNSNSMSLDYVRRGTIVLH |
| | | rna-SPHA_69492 | *Sepia esculenta* | MLPGVGVFGTTATIQSFVPILKVCGFPVVALWGSENLDAQELATQLNIPFSTRKVDDVLLRKDVDLVVIGCPPHSQWFIAVKALGIGKHVLCSAPSGPMQLNAQHMVQAARYYPRLMSLMCYGLRFLPTIVKMKKMIEEGYLGNITICEVKVHYGGLPKEKYDWMCDEGMGGGVLNTIGSNIIDVITFLTKERAVRVHGMLKTYTKQTQNIKGIREITSDDFCTFQMELDKGACVTVTLNSHIPGQFVQEIVICGQKGRLIARGSDLYEQRLNATRESLIHFDPIKEEERYGISPKARTEIPSPYLKGLIHMIEAVKDAFEKEEERQNWQAEPVASASNFEDALYVQTVVDAIRKSNKTKQWTKVDVSVEEPEPNSNMLSDQMRRSTFSLQ |
| | | rna-SPHA_69492 | *Sepia officinalis* | MLPGVGVFGTTATIQSFVPILKVCGFPVVALWGSENLDAQELATQLDIPFSTRKVDDVLLRKDVDLVVIGCPPHSQWFIAVKALGIGKHVLCSAPSGPMQLNAQHMVQAARYYPRLMSLMCYGLRFLPTIVKMKKMIEEGYLGNITICEVKVHYGGLPKEKYDWMCDEGMGGGVLNTIGSNIIDVITFLTKERAVRVHGMLKTYTKQTQNIKGIREITSDDFCTFQMELDKGACVTVTLNSHIPGQFVQEIVICGQKGRLIARGSDLYEQRLNATRESLIHFDPIKEEERYGISPKARAEIPSPYLKGLIHMIEAVKDAFEKEEERQNWQAEPVASASNFEDALYVQTVVDAIRKSNKTKQWTKVDVSVEEPEPNSNMLSDQMRRSTFSLQ |
| chrZ (sex) | OG0008779 | pred2_09431.1 | *Architeuthis dux* | MNGELKEKFEPHKSSDFLERISQKSISEDIKTEIQPTGNLKFRSEQLAYTRSGDRRNTADIAPPFQPSQKSSPAGSFKDFHKNVDMFSQGDNKIHIHGMTRDCHQAPFD |
| | | Dopeav2130673m.g | *Doryteuthis pealeii* | MKLLSNICLHLCRNKQYLFRRNLLSVVPMTRRHNHQENGKRPSSVRIGCGSGFWGDSSIAAQQLIHYGKIDFLVFDYLSEITMSLLTAMKQKNPDMGYAPDFVHFSMVPHLGTIKEKGIKVVSNGGGVNPHACAKLLGDFCKKQNVDLNIAVVTGDDLMIELKKIKQLNLKDMSSGMEFPKTVHSMNAYLGAGPIARALDLGADIVVTGRCVDSALVLGPLLHTFKYKKNDFDQLASGSLAGHIVECGAQATGGNFSDWHTVQNWDTIGFPIVEFSDDGSFIVTKPQSSGGCVTRATVCEQMLYEIGDPKRYQLPDVTADFSQVQINEIEGMTDAVLVKGAKGTPPSDSYKITATYADGFRATAVACVGGPRSEEKAIKTADAILERCRKMFKMLNLGDFTKVHVEILGSEPNSSGSQIIPRQLGLWLAVHHSNKKALEFFAREIAPAGTGMAPGLTGIVGGRPRVSPVLKLFSFLYPKDNLNVEIFINGEQKEKFEPFKNSDIPQTIKSEEPTTEDAKSENLPVGNFNFRLEDLAYTRSGDKGNSANIGVIARDPSYLPYLRQALTEEAMEQYFSYLFEDNNGVRVTRYDVPGIDGINFVLHNSLGGGGIASLRSDPQVRKENIYYHYLEEKCNSQGLVSDL |
| | | g22135.t1 | *Euprymna scolopes* | MTTRLNHHGKPLKSVRIGCGSGFWGDSSIAAEQLIHQGKIDFLVFDYLSEITMSLLTAVKNKHPKLGYTPDFVQYSLLPHLKAIKKKGIKVISNGGGVNPHACGNLIMDFCKKNDIDLNVAVVTGDDLMSELKKMEQFGPKEMLSGMEFPKTVNSMNAYLGAGPIARALDLGADIVVTGRCVDSALVVGPLLHKFKYKMSDFDLLASASVAGHIVECGAQATGGNHSDWKIVKDWANIGFPIVDFSKDGSFIITKPDSTGGCVTCGTVGEQMLYEIDDPQKYLLPDVTVDFSRIQIREIEGRTDAIQVVGAKGYPPTKDYKVTATYADGYRATAVACVGGPRSEEKARKTANALLERCRSWFKRLNLGDFSKVNIEILGSETEQPFSQVVPRQLALWLAVNHPKKNALEFFAREIASAGTGMAPGLTGIVGGRPRISPVLKLFSFLYPKDSINVEIFINGEMKEIFYQDLDLCSDSSEIVKPVCEDIHTETLLTGNFNYTLQDLAYTRSGDKGNSANIGVIARDPSYLPYIKRALTAEAVKEFFMYAFEDTEKARVTRYNVPGIHGLNFVLHDSLGGGGIASLRSDPQGKAYGQILMDFPIKNVPKLLP |
| | | LOC_00003890-mRNA-1 | *Illex illecebrosus* | MRLLKSTCLHLHRSRQCLFSSRNLLSVVPMTKRYNNLQVSSLSGERPNKAVRIGCASGFWGDSSVAAQQLIHHGKIDYLVFDYLSEITMSLLTAAKQRNPDMGYAPDFIHFSVAPHLQMIKDKGIKVVSNGGGVNPHACAKLLKDFCEKAKIDLNIAVVTGDDLMPEIKKIKELSPKDMSSGMEFPKAVHSMNTYLGAGPIARALDLGADIVVTGRCVDSALVLGPLLHTFKYKNNDFDQLAAGSLAGHIVECGAQATGGNFSDWYTVENWDSIGFPIVEFSEEGSFIVTKPESTGGCVTRATVSEQMLYEIGDPKRYLLPDVTADFSQVHIKEIEGMSDAVLVQGAKGTPPSNSYKVTATYADGFRATAVACIGGPRSEEKGLKTANSIFEKCRKMFKKMNMADFSKVHVEILGSESQTSPSQSTPRQVGLWMAVHHYNKKALEFFAREIAPAGTGMAPGLTGIVGGRPRVSPVLKLFSFLYPKDDINVEIFMNGELKERFEQFETTEFPETVDQKPTEDVKNEKLPRGNFQFRLEDLAYTRSGDKGNSANIGVIARDPSYLPYLREALTEEAVREYFSYLFEDSKSGRVTRYDVPGVHGLNFVLHDSLGGGGIASLRSDPQGKAFGQILMDFPITGVPNLCNKK |
| | | obimac_0008956.1 | *Octopus bimaculoides* | MSASRMSSKLFLNNRYSLKKFVSFNRQYCYKIPNEHDVVRIGCASGFWGDTAVAAPQLIHHGKIDFLVFDYLSEITMSLLTAAKQKNPAMGYAPDFIHFSLAPYLKTIKEKKIRVVSNAGGINPLSCAEVLKKLSKEAGVDMNIAVVTGDDLMPKVKEISQSNIQEMSSGMKIPKTIHSMNAYLGAGPIARALDLGADIVVTGRCVDSALVLGPLIHAFKYNNSNFDQLASASLAGHIIECGAQATGGVFTDWQTVHNWDNIGFPIVEFSANGSFSVSKPPSTGGLVSKGTVCEQLLYEIGDPKNYILPDIICDFSQVQVFESEKDSVQVKGALGKPPTNDYKVTATYADGYRVTAVTCIGGPQAEEKAKKTTDAILKRCRNIFKQLKLGDFIRVNVEILGFESKNENKEEPRQLAVWMAAHHPKKQALEILSREIAPAGTGMAPGLTAIAGGRPRVSPVLKLFSFLYPKDELKVQIYMNGELKESYQPSEYSPTLKENVSESSANSEESNLPTGNCEFYLKDLAYTRSGDKGNSANIGVIARHPSYLPYLRAALTESAVEKYFSSLFEDDDDGKRVTRYDVPGINAMNFVLHNCLGGGGIASLQSDPQGKALGQHLLNFKITNVPDLLSKIF |
| | | XP_029648810.2 | *Octopus sinensis* | MSASRMSAKLLLNNRYSLKKFVCFNRQYCYKIPNENDVVRIGCASGFWGDTAVAAPQLIHHGKIDFLVFDYLSEITMSLLTAAKQKNPAMGYAPDFIHFSLAPYLKTIKEKKIRVVSNAGGINPLSCAEVLKKLSQEAGVDMNIAVVTGDNLMPKVKEISQSNIQEMSSGMKIPKTIHSMNAYLGAGPIARALDLGADIVVTGRCVDSALVLGPLIHAFKYNNSNFDQLASASLAGHIIECGAQATGGVFTDWQTVHNWDNIGFPIVEFSANGSFSVSKPPSTGGLVSKGTVCEQLLYEIGDPKNYILPDIICDFSQVQVSESGKDNVQVKGALGKPPTNDYKVTATYADGYRVTAVTCIGGPQAEEKAKKTMDAILKRCRNIFKQLKLGDFTRVNIEILGSESKNENNKEPRQLAVWMAAHHPKKQALEILSREIAPAGTGMAPGLTAIAGGRPRVSPVLKLFSFLYPKDELQVQIYMNGELKESYQPSEYSPTSKENVSESSANSEESNLPTGNCEFYLKDLAYTRSGDKGNSVNIGVVARHPSYLPYLRAALTESAVEKYFSFLFEDDDDGERVTRYDVPGINAMNFVLHNCLGGGGIASLRSDPQGKENLINRSIERLIRCNHRFKYTFVQYNRND |
| | | rna-SPHA_78642 | *Sepia esculenta* | MNTLVLIVDDERSGRTIVVVALAVDNVCLSATETAVASSSFSAIAALGEYKFIFFFQLLPKKSKEQ*DY*KIFVYV*TGTGTAWKWEISQNCENWMRKWFLG*QLCCGSATYPSWQN*FSCF*LFI*NHNVSSHCC*TKES*YGLCTRLCTFFFGTLP*NYKR*RYQGCQQCWRCESTCLCQITKRVLHEKEC*PQNCCSNRR*FDA*D*ENETIIS*RHVFWYGIS*NCS*YECLSWCRSY*KGS*PWC*YCCDWQMCGQCFGPWTAPSYFQIQKE*F*PTCIW*SCWSHSRVRSSSYWR*L**LAYS*KLGYNWFPNCGIFRGWLFPCYQTPINWWLCHKSHCL*TNVI*NWRS*EISIT*RDS*F*PSADKGN*RNEGYGPSTRC*RNSTIQ*L*GYCNLC*WFSSYSSCMCWGTSQ*RESNKNS*CYFRKMQKNIQKAKFG*FY*SPCGTLGIRTIYILPSDNSTTIRTLVSSST***NSPGDFCSRDCTSWNRNGTWINRNSWRKTSSISCTEAFFIFVS*RQYHRKSFHKW*AKRKV*VTEKFRIF*NSKPEDNF**QED*NTSRWKLYVQIGRACLHSKW**RKYSKYRGHSTRSFLSSLPQESSYRGSHQRILQLFV*R*QKSTSDKV*CTRYTWNKFCFTQLTWWGWNCISQE*SSGKSPRPDSDGFSNYMCS*PV**V |

| | | | | |
|---|---|---|---|---|
| | | rna-SPHA_78642 | *Sepia officinalis* | LVLFVDDESGGTTVVVALVVDNVCLSATETAVASSSFSAIAALGEYKFIFFFFFNYCRKK*RTMRLLKNICLRLNWNRHCLKVENVPKL*ELDAEVVFGVTALLRIEKMKQLSPKDMSSGMEFPKTVHSMNVYLGSNTKRMILTYLHLVVLLVT**SVELKLPEVTLVTGIQLRIEGMKDMVLVQGAKGTQPSNDYKVTATYADGFRATAVACVGGPRSEEKAMKTADAILERCRKIFKKLNLADFTKVNVELLGSEPDTTCPQTIPRQLGLWLAVQHKNKTALDFFAREIAPAGTGMAPGLTGIVGGRPRVSPVLKLFSFLYPKDNITGSLHEIRLIFLTSRKLLQRKPSKNTSAICLKITEEHK**VMMYQVYME*ILFYTTHLVGVELLLSGLIL |
| chrZ (sex) | OG0008809 | pred2_10461.1 | *Architeuthis dux* | MTSKSKLIKVVLLGDGGVGKSSLMNRFVSNKFDTQSFHTIGVEFLNKELTLGTDSYTLQIWDTAGQERFKSLRTPFYRGADCCLLTFAVDDKRSFENISMWRKEFAYYADIADDNVSSFPFVIIGNKVDVFERQVDTAEAAEWCEQSGHLPYFETSAKDATNVESAFVAAVKRLREFEEASTTNDLKDSSQGNTVDLSKKRDQTSNSGCCN |
| | | Dopeav2114893m.g | *Doryteuthis pealeii* | MMSSKSKLIKVVLLGDGGVGKSSLMNRFVSNKFDTQSFHTIGVEFLNKELSLGNDSYTLQIWDTAGQERFKSLRTPFYRGADCCLLTFAVDDKRSFENISMWRKEFAYYADIADDNVSSFPFVIIGNKVDVSDRQVTTGEAAEWCEQSGNLPYFETSAKDATNVEAAFVAAVKRLREFEEASTNNDLKDSTQGNTVDLSKKRDQISNTGCCN* |
| | | g16856.t1 | *Euprymna scolopes* | MSTKSKLIKVVLLGDGGVGKSSLMNRFVSNKFDTQSFHTIGVEFLNKEITLGNDTYTLQIWDTAGQERFKSLRTPFYRGADCCLLTFAVDDKRSFENISMWRKEFAYYADIADDNVGTYPFVIIGNKIDVSERQVTTAESSEWCEQNGKLPYFETSAKDATNVEAAFVAAVKRLRDFEEASTTNDLKDSSQGNTVDLSKKRDQVNSSGGCCN |
| | | LOC_00002564-mRNA-1 | *Illex illecebrosus* | MTSKSKLIKVVLLGDGGVGKSSLMNRFVSNKFDTQSFHTIGVEFLNKELTLGNDSYTLQIWDTAGQERFKSLRTPFYRGADCCLLTFAVDDKRSFENISMWRKEFAYYADIADDNVSSYPFVIIGNKVDVVERRVDTAEAAEWCEQNGKLPYFETSAKDATNVESAFVAAVKRLREFEEASTTNDLKDSSQGNTVDLSKKRDQSSSTGCCN |
| | | obimac_0008634.1 | *Octopus bimaculoides* | MMGSKSKLLKVVLLGDGGVGKSSLMNRFVSNKFDTQSFHTIGVEFLNKEISIGSESYTMQIWDTAGQERFKSLRTPFYRGADCCLLTYAVDDVKSFENVSMWKKEFLYYADIDDESTFPFVVIGNKIDVGQRLVSYNTASEWCEKNGHVPYFETSAKDSTNVEVAFTAAVKRLKDLEDKCAEVKPSHGNTVDLNKKKESQSTGCCN |
| | | XP_036367568.1 | *Octopus sinensis* | MQAEKLVQQNLVAFILKHLFGCVYEHKQMGSKSKLLKVVLLGDGGVGKSSLMNRFVSNKFDTQSFHTIGVEFLNKEISIGSESYTMQIWDTAGQERFKSLRTPFYRGADCCLLTYAVDDVKSFENVSMWKKEFLYYADIDDESTFPFVVIGNKIDVGQRLVSYNTASEWCEKNGHVPYFETSAKDSTNVEVAFTAAVKRLKDLEDKCAEVKPSHGNTVDLNKKKESQSTGCCN |
| | | rna-SPHA_58078 | *Sepia esculenta* | MTYLMTSLIPPFWNANFYVILLNRFDEKGRFPFNTMSSKSKLIKVVLLGDGGVGKSSLMNRFVSNKFDTQSFHTIGVEFLNKELTLGNDSYTLQIWDTAGQERFKSLRTPFYRGADCCLLTFAVDDKRSFENISMWRKEFAYYADIADDNVSSFPFVIIGNKVDVFERQVTTGEATEWCDQNGKLPYFETSAKDATNVEAAFVAAVKRLREFEEAGTTHDLKDSTQGNTVDLSKKRDQISNSGCCN |
| | | rna-SPHA_58078 | *Sepia officinalis* | MTSLMTSLMPPFWNANFYVILLNRFDEKGRFPFNTMSSKSKLIKVVLLGDGGVGKSSLMNRFVSNKFDTQSFHTIGVEFLNKELSLGNDSYTLQIWDTAGQERFKSLRTPFYRGADCCLLTFAVDDKRSFENISMWRKEFAYYADIADDNVSSFPFVIIGNKVDVSERQVTTGEATEWCDQSGKLPYFETSAKDATNVEAAFVAAVKRLREFEEAGTTHDLKDSTQGNTVDLSKKRDQISNSGCCN |
| chrZ (sex) | OG0009225 | pred2_26014.1 | *Architeuthis dux* | MLQQQPNSLEPVSHFATKAAMPSLSPEWRFLPELAPTNIESNNKESGKESYFPDAGGIKKYNREFLLELRHTRASLLMPQCLPDLPKELLKMPYPSKMFNLSCSDPFCKGFKDRRPAEKSRSDEGKEVNCLTDTLCNKTTERYKPLLSEIRCIQIDTESQLKAIIDLLFEKASNYPILGISYAYLCRSLSLIRVPSATRQGETVNFNKLLNRRCQIELEKIKQEDEAICQTKQMISAENDEARIMKLQMKLGELTSNSNKRTLGNMKLIGEFVKLRILKESIIIQFVCSLLSSRTEARIECLCVLLKTVGMELEHNSNRQDKKKIEDCFTEMKKMVSQGTCSPRVKAILSSIIRLRENKWVF |
| | | Dopeav2060590m.g | *Doryteuthis pealeii* | MLHQQPNSLESVNNFTTKAVISNLSPEWKFSPELATTNMESNNNNKETGQESCFPDAGGIKKYNREFLLQLRHTRASLLMPPCLPDLPKELLKMPYSSKMFNLSCSDPFYKGLKDRSPVEKSRSDESKESNCLMDTFCKIRCIQIDTESQLKAIIDLLFEKASNYPILGISYAYLCRSLSLIRVPSATRQGETVNFNKLLNRRCQIELEKIKQEDEAINQTKQMVRSAESDEPRIIKLQKKLGELTSASKKRTLGNMKLIGEFFKLRILKENIIIQFVCSLVSSRSEDRIECLCVLLKSVGLELDRSSNRQDKKKMEDCYTEMKKIISQGTCSPRVKAMLSSIIRLRENKWVF* |
| | | cluster_18938 | *Euprymna scolopes* | MMSSLAPEWRFLSKSAPTNIEEPRQESCFQSATGIKKYSREFLLQLRHTRASLLMPACLPDLPRELIKMQYSSETFNLSCSDPFFKSLKDRRPSEESRNDKNNETKKPLLSEIRCIQFDTESQLKAIIDLLFEKASKNPILGISYAYLCRSLSLVRVPSSTRQGETVNFNKLLNRRCQIELEKIKQANEAINQAKQMIRSAENGEARDIQLQKKLDELMSTSRKRTMGIMKMIGEFFKMRILNENIITEFVCSLLSPRTEDRVECLCDLLKTVGKELEQNSNRQEKKKMEDCFSEIKKTVSQAGEVHQANYNTTTGLQRIPGGPPPLCQKQLIACADVIRRPSTFARLHRLTGTNQQNEQKES |
| | | LOC_00011915-mRNA-1 | *Illex illecebrosus* | MLQQQPNPLESVSSFTTKAAMPSLSPEWRFSPESAPTNIEPNNKESEKESYFPDAGGIKKYNREFLLQLRHTRASLLMPPCLPDLPRELLKIPYPSKMFNLSCSDPFFKGLKDRRPAEKSRSDEGKEVNCLTDTICSKTTERYKPLLSEIRCIQIDTESQLKAIIDLLFEKASNYPILGISYAYLCRSLSLIRVPSATRQGETVNFNKLLNRRCQIELEKIKQEDEAISQTRQFIRNAENDEARVMKLQKKLAELTTSSKKRTLGNMKLIGEFFKLRILKENIVIQFVCGLLSSRTEDRIECLCILLKTVGTELERNSNRQDKKKMEDCFTEMKKVVSQGTCSPRVKTMLSSIIRLRENKWVF |
| | | obimac_0009088.1 | *Octopus bimaculoides* | MPFSNSPSENAYSYPNSNVSSTEETQEATQRTPLPSQIRLKYDRDFILKLQYKEASLTKPVGLPDLPEILFKKPQIPKQLKYNKIRRDIIPFKCSKNMKSVDEYVNDSEDEELNTKMKNIASRLAPEKYKSVLSQIRDIQIDKESKLVAIMNLLFEKATSDPMLSTAYAYVCRCLTLKRVPSIHCHKEVVNVHKVLNKKCQKEFEKYQVEEAMLNKKLQQIEDTDNNIEKLKFQKELVKDIEIFKRKSQANVRFLCELFKLGVLRENTMQKYIKKMLQSLSENSLESVCLLFNTIGERLDTGKKFKMDVYFIQLKSIAEENSTPNRLKVMIENLISLRENQWNQKEAENHTKQKGMTSVKDSMKEETLKVMANSYYENKVKDIYQNGDGSVASNMGDIFSSDPGTKTSSSYQQPQQQQQQQQHIKEEYVKPGYMKIFDIDAQNKCMDLTQLERKYCNLVYDGTTRASARGDVMVPNCSPNIIRETKEAV |
| | | XP_036367469.1 | *Octopus sinensis* | MPFSNSPSENAYSYPNSNVSSTEETQEATQRTPLPSQIRLKYDRDFILKLQYKEASLTKPVGLPDLPEILFKKPQIPKQLKYNKIRRDIIPFKCSKNMKSVDEYVNDSEDEELNTKMKNIASRLAPEKYKSVLSQIRDIQIDKESKLVAIMNLLFEKATSDPMLSTAYAYVCRCLTLKRVPSIHCHKEVVNVHKVLNKKCQKEFEKYQVEEAMLNKKLQQIEDTDNNIEKLKFQKELVKDIEIFKRKSQANVRFLCELFKLGVLRENTMQKYIKKMLQSLSENSLESVCLLFNTIGERLDTGKKFKMDVYFIQLKSIAEENSTPNRLKLMIENLISLRENQWNQKEAENHTKQKGITSVKDSMKEETLKVMANSYYENKVKDIYQNGDGSVASNMGDIFSSDPGTKTSSSYQQPQQQQQQQQHIKEEYVKPGYMKIFDIDAQNKCMDLTQLERKYCNLVYDGTTRASARGDVMVPNCSPNIIRETKEAV |

| | | | | |
|---|---|---|---|---|
| | | rna-SPHA_31329 | *Sepia esculenta* | MLHQQPNSIEPVTSFATKAAMPSLSPEWKFSPESAPTNIESANRESGPESCFPDAGGIKKYNREFLLQLRHTRASLLMPSCLPDLPRELLKMPYSSKTFNLSCSDPFFKGLKDRRPAEKSRSDESKEGNCFTDSSKMAERYKPLLSEIRCIQIDTESQLKAIIDLLFEKASNYPILGISYAYLCRSLSLIRVPSATRQGETVNFNKLLNRRCQIELEKIKQEDETINQTKQMIRSAEVDETRIIKLQKKLCELTSTSKKRTLGNMKLIGEFFKLHILKENIVIQFVCNLLSTRTEDRIECLCVLLKTVGMELERNSNRQEKKKLEDCFTEMKKIVSQGTCSPRVKSMLSSIIRLRENKWVF |
| | | rna-SPHA_31329 | *Sepia officinalis* | MLHQQPNSIEPVTSFATKAATPSLSPEWKFSSESAPTNIESTNRESGPESCFPDPGGIKKYNREFLLQLRHTRASLLMPSCLPDLPRELLKMPYSSKTFNLSCSDPFYKGLKDRRPAEKSRSDESKEGNCFTDSSKMAERYKPLLSEIRCIQIDTESQLKAIIDLLFEKASNYPILGISYAYLCRSLSLIRVPSATRQGETVNFNKLLNRRCQIELEKIKQEDETINQTKQMIRSAEVDETRIIKLQKKLCELTSTSKKRTLGNMKLIGEFFKLRILKENIVIQFVCNLLSTRTEDRIECLCVLLKSVGMELERNSNRQEKKKLEDCFTEMKKIVSQGTCSPRVKSMLSSIIRLRENKWAF |
| chrZ (sex) | OG0009585 | pred2_41298.1 | *Architeuthis dux* | MSFLTTDIKALADLLKQEDSDSDGEQQQGSALLGPGHIGGRTKEQETIPSEKDCGQSKEIWNPQEIIEGSEFDSLSDPRPQPEYEILYKQSVTTEDIFLQMGNKTPNTSSCENMVIKIKLPNTELKNITLDVKSIFVDVRTPKYKLGLHLPHPVEEQESNAKWDACQESLIITLKMKREFDFMNY |
| | | Dopeav2130641m.g | *Doryteuthis pealeii* | MGTTKLGSAKFGPGHVGETKKQENISSRKNWALTTQSTVTEENPSAREHSQLHHLSTLSAREINNYFSQSKEIWKPEEIIEGSEFDSLSDPRPQPEYEIFYKQSVTTENIFLQMGNKNPSTSSCENMVIKIKLPNTEMKNITLDIKSVFLDVRTPKYKLGLHLPHPVDEKESNAKWDGCQETLIVTMKMNREFDFLNY* |
| | | g28227.t1 | *Euprymna scolopes* | MSFRVSEIAALTNLLTPANEESDSDEDQPQLGSSKFGPGHIGESKKQRNSASKKDINQSKEIWCSEEITEGAEFDSLSDPRLQPELCHSVHYLQGRHKVTVYQ |
| | | LOC_00003860-mRNA-1 | *Illex illecebrosus* | MSFMTSDLLALTDLLREPEDSDSDEETRYQGTARLGPGHIGKNQKQETSSSKKDISQSKEIWNPEEINQGSEFDSLSDPRPQPEYEILYKQAVTSEDIYLQMGNKTPNTSSCENMVVKIKLPNTEMKNISLDVKSKFLDLRTPKYKLGLHLPHPVAEQESNAKWDSDCSSLIVTLKMNRELDFMNY |
| | | obimac_0008966.1 | *Octopus bimaculoides* | MAFQQCDIRALANLLREPAEDSDSDTDIVCSDYSYGPGHIGPEKNSKDGTAEKDTKQSKDIWSADEIPQGSEFDSLWDQRLQPEYSIVYNQNVRTEDIFLQMGNKTPGSSSCERMVVKIQLPNTMMKDISLDVKPKFLDLRTPKYKLGLHLPFTVKENESQAQWDGGESCLSVSLKMIREYDFVNF |
| | | XP_036367510.1 | *Octopus sinensis* | METIVEIAIMETNEESSTTSNKDEDDFFRNITRFWESRSHRSLKSKVQNLVKTWLEAILKAVLTEVAFFWVCGGHSYGPGHVGPEKNSKDGTAEKDTKQSKDIWSADEIPEGSEFDSLWDQRPQPEYSIVYNQNVRTEDIFLQMGNKTPSSSSCERMVVKIQLPNTMMKDISLDVKPKFLDLRTPKYKLGLHLPFTVKEDESQAQWDGGESCLSVSLKMIREFDFVNF |
| | | rna-SPHA_78646 | *Sepia esculenta* | MVTKITRTSILEAITFGKNIFYRFYKMSFSVADIRSLTNLLKQPDEDSDSDGEQLYQGSASLGPGNIGESEKQEKPTFKKDLSQSKEIWHPEEINAGSEFDSLSDPRPQPEYEILYKQSVTTEDIFLQMGNKTPNTSSCENMIIKIKLPNTEMKNITLDVKSIFLDVRTPKYKLGLHLPNPVKEQESTAQWHSDQESLIVTLKLNREFDFLNY |
| | | rna-SPHA_78646 | *Sepia officinalis* | YEILYKQSVTTEDIFLQMGNKTPNTSSCENMIIKIKLPNTEMKDITLDVKSIFLDVRTPKYKLGLHLPHPVKEQEGAAKWDTDQESLIVNLKLNREFDFMNY* |

**Table S1.**
